# Static and dynamic functional connectomes represent largely similar information

**DOI:** 10.1101/2023.01.24.525348

**Authors:** Andraž Matkovič, Alan Anticevic, John D. Murray, Grega Repovš

**Author notes:** Corresponding author Email address (Andraž Matkovič).

## Abstract

Functional connectivity (FC) of blood-oxygen-level-dependent (BOLD) fMRI time series can be estimated using methods that differ in sensitivity to the temporal order of time points (static vs. dynamic) and the number of regions considered in estimating a single edge (bivariate vs. multivariate). Previous research suggests that dynamic FC explains variability in FC fluctuations and behavior beyond static FC. Our aim was to systematically compare methods on both dimensions. We compared five FC methods: Pearson’s/full correlation (static, bivariate), lagged correlation (dynamic, bivariate), partial correlation (static, multivariate) and multivariate AR model with and without self-connections (dynamic, multivariate). We compared these methods by (i) assessing similarities between FC matrices, (ii) by comparing node centrality measures, and (iii) by comparing the patterns of brain-behavior associations. Although FC estimates did not differ as a function of sensitivity to temporal order, we observed differences between the multivariate and bivariate FC methods. The dynamic FC estimates were highly correlated with the static FC estimates, especially when comparing group-level FC matrices. Similarly, there were high correlations between the patterns of brain-behavior associations obtained using the dynamic and static FC methods. We conclude that the dynamic FC estimates represent information largely similar to that of the static FC.

## 1. Introduction

Brain functional connectivity (FC) is estimated by calculating statistical associations between time series of brain signal [1], which reflect functional relationships between brain regions [2]. The investigation of FC has improved our understanding of brain function in health and disease and has been shown to be useful as a tool to predict interindividual differences, such as cognition, personality, or the presence of mental or neurological disorders [3, 4]. In functional magnetic resonance imaging (fMRI) studies, FC is most commonly estimated using the Pearson’s correlation coefficient between time series of pairs of regions. Although correlation is simple to understand and compute, it is insensitive to the temporal order of time points. Measures or models that are sensitive to the temporal order of time points are called *dynamic*, while measures that are insensitive to temporal order are measures of *static* FC. Given that the information flow in the brain is causally organized in time [5, 6], dynamic connectivity models could be more informative in terms of understanding brain function and investigating brain-behavior associations.

In FC temporal information can be represented in two ways. First, the temporal order of the time points can be taken into account when computing the FC estimates. Models that are sensitive to temporal order are called *dynamic* models, whereas models that do not take temporal order into account are called *static* FC models. Second, the methods can investigate, whether and how FC estimates change over time. A time series model is *stationary* (in a weak sense) if its first- and second-order statistics (mean and variance) do not vary as a function of time [7, 8]. Importantly, the distinction between dynamic and static FC should not be confused with the distinction between stationary and non-stationary FC.

Nonstationarities are commonly estimated using measures of time-varying functional connectivity (TVFC), such as sliding window correlation (SWC) [9, 10]. In this method, we calculate connectivity (e.g. using correlation) in a time window of selected length around a given time point; this window is continuously being moved from the start to the end of the recording. Procedures such as autoregressive (AR) randomization or phase randomization can be used to construct surrogate time series, which can then be used to perform a statistical test of the null hypothesis that a time series is stationary, linear, and Gaussian [11]. Using these procedures Liégeois et al. [7] and Hindriks et al. [12] have shown that the hypothesis that FC is stationary cannot be rejected for most participants. Similarly, Laumann et al. [13] concluded that variation in FC over time within a single session can be largely explained by sampling variability, head motion, and fluctuating sleep state. Furthermore, EEG FC has been shown to be largely stable during resting state [14], during sleep [15], and even before, during, and after epileptic seizures [16]. On the other hand, several studies rejected the stationarity hypothesis for certain connections [17, 18, 19]. However, do note that Zalesky et al. [19] found that on average only 4% of the connections are nonstationary.

The inability to reject the stationarity hypothesis does not imply the absence of brain states [7], nor does it preclude finding (behaviorally) relevant information using models of TVFC [11]. However, if a simpler model (i.e., a more interpretable model with fewer parameters) can be used to describe FC dynamics, it should be preferred to a complex model (such as SWC) unless the simpler model cannot model some important aspect of the time series a researcher is interested in [7, 20]. Indeed, recent work on resting state fMRI FC in humans has shown that many of the properties of TVFC can be predicted from a static and stationary FC model [20, 21, 13, 22]. Similarly, Liégeois et al. [7] showed that SWC fluctuations can be explained with a model of dynamic (and stationary) FC. Since many studies have shown that FC is largely stationary, we focused on the relationship between *dynamic* (as defined above) and *static* connectivity.

Dynamic FC can be estimated using measures of lag-based connectivity, such as lagged correlation or multivariate autoregressive (AR) model. In contrast to static FC, dynamic FC methods can be used to estimate the *directionality* of information flow based on temporal precedence [23]. Although these methods have been commonly used, some studies [23, 24, 25, 26, 27] have warned that the ability of these methods to accurately estimate the presence and directionality of connections is compromised due to the convolution of the neural signal with the hemodynamic response function (HRF) and the resulting blurring of the signal, due to interregional variability of HRF [26, 24, 25], noise [23, 24, 26], and/or downsampling of the neural signal in fMRI [27]. Other studies [28, 29, 30, 31] have shown that the measures of dynamic FC complement the measures of static FC. For example, lagged FC measures can improve discrimination between individuals and between tasks [28, 29], have better predictive value for PTSD compared to static FC [31], and can be used to improve effective connectivity estimates [30]. Furthermore, Liégeois et al. [7] have shown that the multivariate AR model explains temporal FC fluctuations better than Pearson correlations.

In subsequent research Liégeois et al. [32] showed that static FC and dynamic FC exhibit different patterns of brain-behavior associations. They concluded that dynamic FC explains additional variance in behavior beyond variance that can be explained by static FC. However, this comparison confounds two orthogonal properties of FC methods. Although Pearson’s correlation and multivariate AR models differ in their sensitivity to temporal reordering (i.e., static vs. dynamic), they also differ in terms of how many variables (brain regions) are taken into account during the estimation of a single edge (bivariate vs. multivariate). Hence, a more valid comparison between static and dynamic FC methods should consider both dimensions: the number of variables and the sensitivity to temporal reordering. Combining these two factors enabled us to differentiate between four basic classes of FC methods (see Figure 1B).

**Figure 1:**
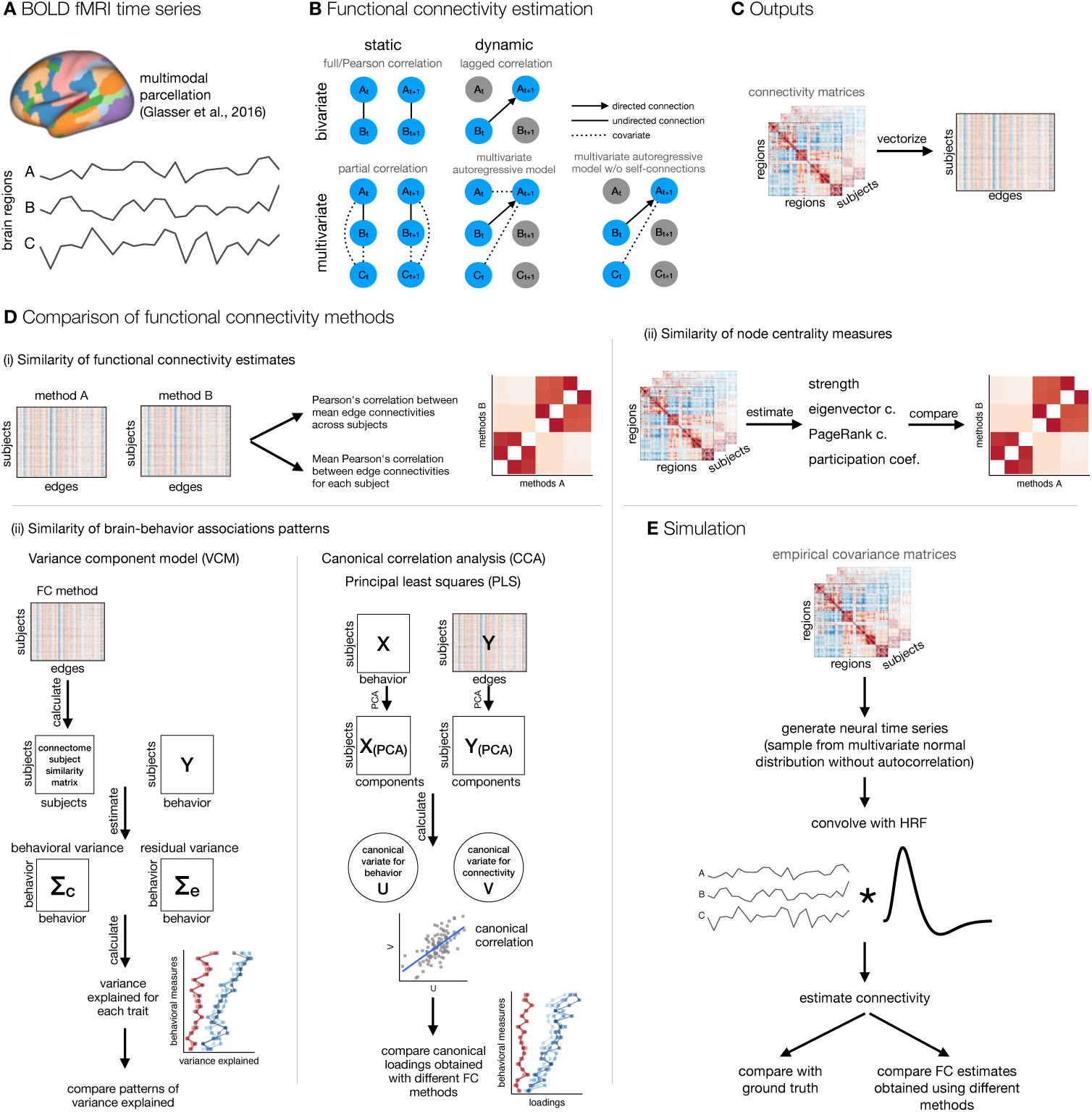
A schematic of analysis steps. **A.** BOLD fMRI data was preprocessed, parcellated, and individual parcel timeseries were extracted. **B.** Functional connectivity (FC) was estimated with five methods that differed along two dimensions: static vs. dynamic and bivariate vs. multivariate. Static FC refers to measures that are insensitive to temporal order and can be estimated using full/Pearson’s correlation or partial correlation, whereas measures of dynamic FC are sensitive to temporal order of time points. Dynamic FC can be estimated using measures of lag-based connectivity, such as lagged correlation, or using the linear multivariate autoregressive (AR) model. The lagged correlation between two time series is calculated by shifting one time series by *p* time points. Similarly, a *p*-th order multivariate (or vector) autoregressive model predicts the activity of a particular brain region at time point *t* based on the activity of all regions at time point(s) from *t p* to *t* 1. Bivariate and multivariate FC methods differ in terms of number of variables (regions) taken into account when estimating connectivity at a single edge: bivariate connectivity between two regions depends only on the two regions, whereas multivariate connectivity between two regions includes all other regions as covariates. **C.** FC matrices were vectorized. **D.** FC estimates were compared (i) by calculating correlations between FC estimates, (ii) by calculating correlations between node centrality measures, and (iii) by comparing estimates of brain-behavior associations across FC methods. **E.** Additionally, we performed simulation to assess the influence of random noise and signal length on the similarity between FC estimates obtained using different methods.

Our aim was to systematically compare the FC estimated by both dimensions, that is, the sensitivity to temporal reordering (static vs. dynamic) and the number of independent variables (bivariate vs. multivariate). We focused on five mathematically related methods: full/Pearson’s correlation, partial correlation, lagged correlation, and multivariate AR model with and without self-connections, where self-connections refer to autocorrelation of the region with itself [33, 34]. We were interested in similarities of the FC estimates and patterns of brain-behavior associations. We compared FC methods (i) by assessing similarities between FC matrices, (ii) by comparing node centrality measures, and (iii) by comparing brain-behavior associations. In addition, to better understand the results obtained using different methods and the relationship between them, we generated and analyzed synthetic data in which we systematically varied the length of time series and the amount of noise.

We used empirical and simulated data to test several hypotheses. First, we predicted that dynamic and static FC methods will provide similar FC estimates due to autocorrelation of the fMRI time series. Autocorrelation is inherent to the fMRI signal and originates from two main sources: physiological noise and convolution of neural activity with HRF [35]. We expected the degree of similarity between static and dynamic FC estimates to be similar to or greater than the average autocorrelation of the fMRI time series. Furthermore, we expected the similarity between dynamic and static FC to be lower when the fMRI time series is pre-whitened (i.e., when autocorrelation is removed before computation of FC).

Second, we predicted that multivariate methods can improve inferences about causal relationships between regions, as they estimate *direct* connections by removing the confounding influence of indirect associations [2] as opposed to bivariate methods, which cannot separate *indirect* and *direct* connections [34]. By providing more direct information on causal relationships between brain regions [36], multivariate methods could improve brain-behavior associations in terms of explained variance and/or brain-behavior correlation estimates. Existing research has shown inconsistent differences in behavior predictive accuracy between partial and full/Pearson’s correlations, favoring either partial [37, 38] or full correlation [39] or showing negligible differences between them [40].

Finally, the choice of FC method can affect the measures of network topology [e.g. 41, 42]. To address this problem, we compared FC estimates using common node centrality measures, including strength, eigenvector centrality, PageRank centrality, and participation coefficient. Using centrality measures also allowed us to compare incoming and outgoing connections in dynamic FC estimates. Based on previous research showing that nodes are either receptors or feeders (i.e., they have predominantly incoming or outgoing connections), but not both [30], we expected a negative correlation between in-degree and out-degree. We had no specific hypotheses regarding the similarity of node centrality measures between multivariate and bivariate methods or between static and dynamic FC methods.

## 2. Method

### 2.1. Participants

To address the research questions, the analyzes were performed on publicly available deidentified data from 1096 participants (*M_age_* = 28.8, *S D_age_* = 3.7, 596 women) included in the Human Connectome Project, 1200 Subjects Release [43]. Each participant took part in two imaging sessions over two consecutive days that included the acquisition of structural, functional (rest and task), and diffusion-weighted MR images. The study was approved by the Washington University institutional review board and informed consent was signed by each participant.

### 2.2. fMRI data acquisition and preprocessing

Data were acquired in two sessions using the Siemens 3T Connectome Skyra tomograph. Structural MPRAGE T1w image (TR = 2400 ms, TE = 2.14 ms, TI = 1000 ms, voxel size = 0.7 mm isotropic, SENSE factor = 2, flip angle = 8^◦^) and T2w image (TR = 3200 ms, TE = 565 ms, voxel size = 0.7 mm isotropic) were acquired in the first session. The participants underwent four resting state fMRI runs, two in each session (gradient echo EPI sequence, multiband factor: 8, acquisition time: 14 min 24 s, TR = 720 ms, TE = 33.1 ms, flip angle = 52^◦^).

Initial preprocessing was performed by the HCP team and included minimal preprocessing [44], ICA-FIX denoising [45] and MSMAll registration [46]. The data was then further processed using QuNex [47] to prepare them for functional connectivity analyzes. First, we identified frames with excessive movement and/or frame-to-frame signal changes. We marked any frame that was characterized by frame displacement greater than 0.3 mm or for which the frame-to-frame change in signal, computed as intensity normalized root mean squared difference (DVARS) across all voxels, exceeded 1.2 times the DVARS median across the time series, as well as one frame before and two frames after them. Marked frames were used for motion censoring, which is described in detail below. Next, we used linear regression to remove multiple nuisance signals, including six movement correction parameters and their squared values, signals from the ventricles, white matter and the whole brain, as well as the first derivatives of the listed signals. The previously marked frames were excluded from the regression and all subsequent analysis steps were performed on the residual signal. No temporal filtering was applied to the data, except a very gentle high-pass filter at the cutoff of 2000 s applied by the HCP team [44], since temporal filtering could introduce additional autocorrelation [48] and inflate correlation estimates [35, 49]. Motion scrubbing is usually performed by removing frames thought to be affected by movement (i.e. *bad* frames) before calculating the correlation or related measure of static FC. This is not appropriate in the case of dynamic FC or autocorrelation, since removing time points disrupts the autocorrelation structure of time series. To overcome this limitation, a frame was considered bad if it was bad in either original or lagged time series. Frames at transition between concatenated time series (last frame in the first time series and first frame in the next time series) were also marked as bad in this case.

Only sessions with at least 50% useful frames after motion censoring were used in the further analysis, except where noted otherwise. This resulted in 1003 participants with at least one session. Before FC analyzes, all resting-state BOLD runs from available sessions were concatenated and parcellated using a multimodal cortical parcellation (MMP1.0) containing 360 regions [50]. Each parcel was represented by a mean signal across all the parcel grayordinates.

### 2.3. Functional connectivity estimation

Functional connectivity was estimated using five methods: full (Pearson’s) correlation, partial correlation, lagged correlation, multivariate AR model (also called vector AR model), and multivariate AR model without self-connections. The listed methods differ in terms of the number of regions used to estimate the connectivity of a single edge (bivariate vs. multivariate) and in terms of sensitivity to temporal reordering of time points (static vs. dynamic) (see Figure 1B). A multivariate AR model without self-connections was included to test how much similarity between the multivariate AR model and partial correlation depends on self-connections (the diagonal terms in the autocovariance matrix).

The bivariate static FC was estimated using full correlation. Let *x_i_* be a demeaned *T* × 1 vector of region *i* time series (*T* is the number of time points) and let *X* = [*x*_1_, . . ., *x_N_*]^′^ be a *N T* matrix of the demeaned region time series (*N* is the number of regions). Then the sample covariance matrix *C* can be estimated with

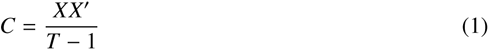

A correlation matrix can be obtained by standardizing the time series to zero mean and unit standard deviation (i.e., *z*-scores) beforehand.

Multivariate static FC was estimated using partial correlation. Partial correlations were computed by taking an inverse of a covariance matrix (i.e., the precision matrix) and then standardizing and sign-flipping according to the equation:

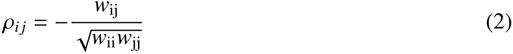

where *ρ* is an element of a partial correlation matrix, *w* is an element of a precision matrix, and *i* and *j* are the indices of rows and columns, respectively [51].

Dynamic bivariate connectivity was estimated using lagged correlation (also known as autocovariance matrix). Autocovariance is defined as the covariance of time series with lagged time series. Let *X_t_* be an *N* (*T p*) matrix of shortened time series with time points from 1 to *T p* (*p* is the lag/model order) and *X_t_*_+*p*_ be a similar matrix with time points from *p* + 1 to *T* . Then,

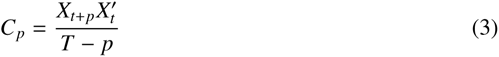

is *p*-th order autocovariance or lagged covariance matrix. Diagonal entries are called autocovariances or, sometimes, self-connections or self-loops [34, 33]. Off-diagonal entries of autocovariance matrix are also called cross-covariances. Note that the autocovariance matrix of lag 0 is equal to the ordinary covariance matrix. The autocorrelation matrix was obtained by standardizing time series before computing autocovariance.

Correlations, autocorrelations, and partial correlations were Fisher *z*-transformed for subsequent analyzes.

Multivariate dynamic connectivity was estimated using the Gaussian multivariate AR model. Let *Z* be an *Np* × (*T* − *p*) matrix of stacked matrices of shortened time series, *Z* = [*X_t_*^′^_+_*_p_*_−1_, . . ., *X_t_*^′^_+1_, *X_t_*^′^]^′^. The multivariate AR model can be written in matrix notation as:

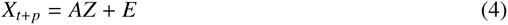

where *A* is an *N × Np* matrix of AR coefficients of the *p*-th order model and *E* is an *N* (*T p*) matrix of zero-mean, independent, normally distributed residuals. The matrix *A* can be estimated using the ordinary least squares (OLS) estimator:

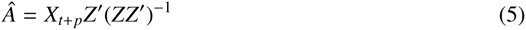

For *p* = 1 *Â* equals:

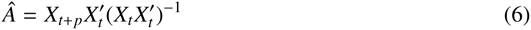

The equation shows that the coefficients of the multivariate AR model are a product of the lagged covariance and (non-lagged) precision matrix. Therefore, the multivariate AR model encodes both static and dynamic FC. The same can be inferred from the Yule-Walker equations [see 7, 8]. Moreover, for lag 0, the coefficients of the multivariate AR model are equal to the covariance matrix [see 7].

To estimate the coefficients of the multivariate AR model without self-connections, we fitted the model

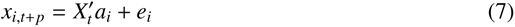

for each region *i* separately, such that we set *i*-th row of matrix *X_t_* to zero (the equation above applies for *p* = 1 only, but the model could be extended to include higher lags as in Equation 4). Vectors *x_i_*_,*t*+*p*_ were taken from rows of the matrix *X_t_*_+*p*_ and included time points from *p* + 1 to *T* . Vectors *e_i_* represent normally distributed, zero-mean, independent residuals. FC matrix was constructed by organizing *N* × 1 vectors *a_i_* into the *N × N* matrix *A*_1_ = [*a_i_*, . . ., *a_N_*]^′^. This matrix is asymmetric with zeros on the diagonal. The coefficients of both multivariate AR models were estimated using the coordinate descent algorithm implemented in the GLMnet package for MATLAB [52].

All AR models were estimated for lag 1 only. This order was shown to be optimal for the multivariate AR model for resting state fMRI data with a high number of regions [53, 54], and also in a study using HCP data [55]. There were no differences between the variance of order 1 and the higher-order models explained by the first principal component of the null data generated from the multivariate AR model in a previous study [7]; therefore, we did not consider higher-order autoregressive models.

### 2.4. Prewhitening

We expected that FC estimates based on AR models would be similar to static FC estimates due to autocorrelation present in the fMRI time series. To test the similarity between static and dynamic FC in the absence of autocorrelation, we computed connectivity from from non- prewhitened time series and prewhitened time series. The exception was the multivariate AR model, where the diagonal terms (self-connections) effectively act as prewhitening. The difference between the multivariate AR model with and without prewhitening is essentially the difference between the multivariate AR model with and without self-connections. We performed the prewhitening by taking the residuals of the regression of the ”raw” time series on the lagged time series.

To retain frame sequence after prewhitening, frames that were marked as bad in any of the original or lagged time series were set to zero before computing residuals. For this reason, frames that were preceded by a bad frame in any of the 1 to *l* previous frames were not prewhitened. At higher orders, this resulted in fewer total prewhitened frames.

Prewhitening was performed on orders 1 to 3 (abbreviated AR1/2/3 prewhitened). Autocorrelations were already significantly reduced at order 1 and were additionally reduced at lags 2 and 3 (Figure 2B). Since the results were similar regardless of the prewhitening order, only the results for the prewhitening on order 1 are shown in the main text, and the results for higher orders are shown in the supplement.

**Figure 2:**
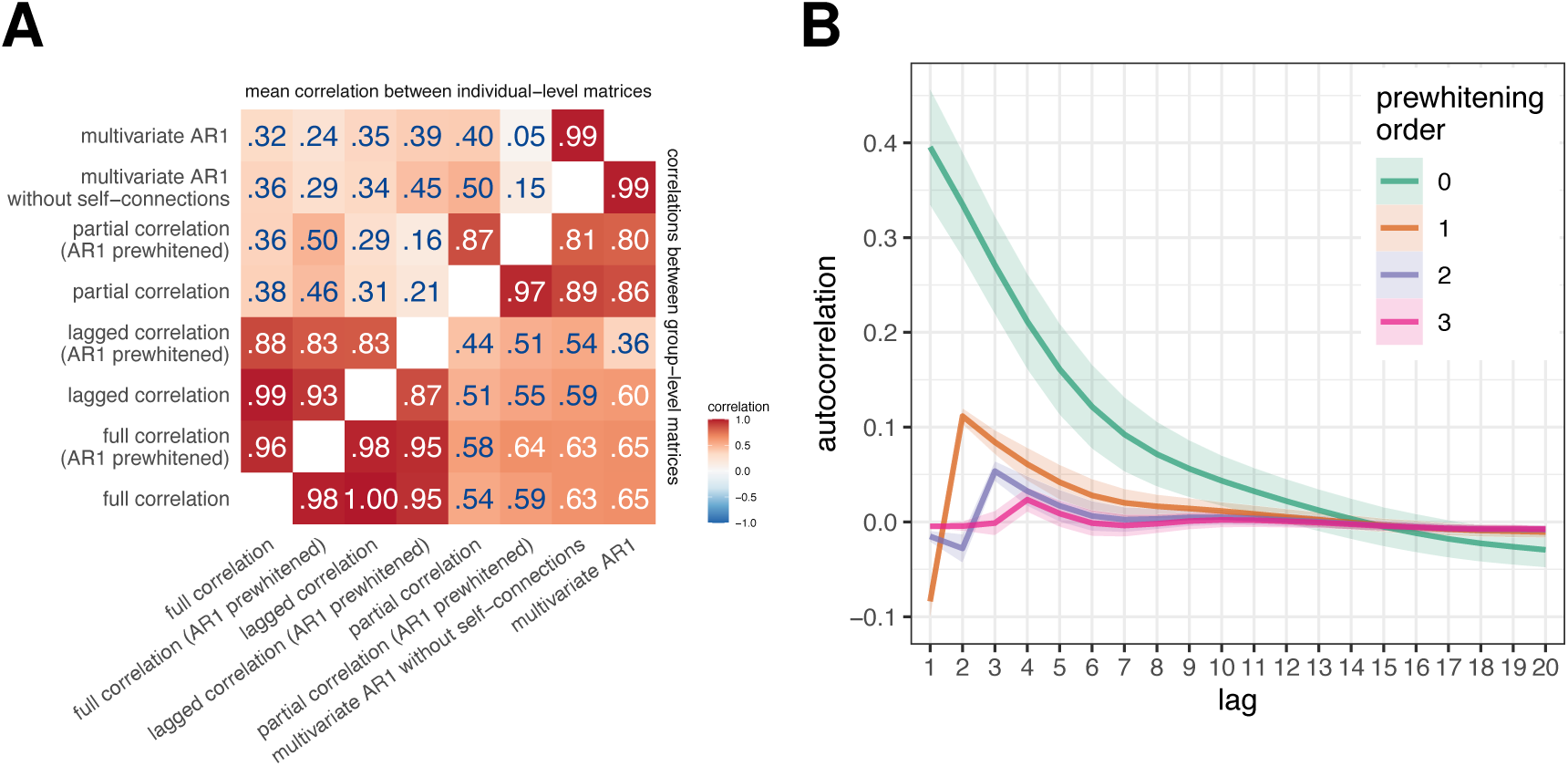
**A.** Correlations between FC estimates obtained using different FC methods. We calculated the similarities between FC estimates obtained using different FC methods (i) by averaging connectivity matrices across participants and then computing correlations between them (correlation between group-level FC, bottom right triangle), and (ii) by computing correlations between the FC estimates for each participant separately and then averaging across participants (correlation between individual-level FC, top left triangle). **B.** Autocorrelation function of experimental data as a function of prewhitening order. The mean autocorrelation function was computed over all participants and regions; the ribbons represent the standard deviation.

### 2.5. Similarities between FC estimates obtained using different methods

We estimated similarities between the FC estimates by computing the correlation between vectorized FC matrices. We adjusted the vectorization for each pair of methods so that only unique elements were taken into account. For example, correlation and partial correlation matrices are symmetric; therefore, only the upper or lower triangular part of the matrix (without the diagonal) should be considered. On the other hand, the FC matrices derived from the AR models are not symmetric; therefore, the whole matrix must be vectorized. The exception is the multivariate AR model without self-connections, which does not contain any information on the diagonal, so in this case matrix without the diagonal needs to be vectorized. When comparing asymmetric and symmetric matrices, we computed and used the average of the upper and lower triangular parts of the matrix (using equation (*X* + *X*^′^)/2).

We estimated similarities in two ways: first, by computing correlations between connectivity estimates for each subject separately and then averaging the resulting correlations (mean correlations between individual-level FC matrices), and second, by averaging FC matrices over participants and then computing correlation between methods on group FC matrices (correlations between group-level FC matrices).

To test how similarity between FC estimates depends on data quality, we repeated analyses on a subset of 200 participants with the largest number of retained frames.

#### 2.5.1. Correlation between edge similarity and test-retest reliability

To better understand the origin of the similarities between the FC methods, we examined the relationship between the edge similarity of the FC estimates obtained using different methods and test-retest reliability at the edge level. If similarities between FC estimates depend on the signal-to-noise ratio (SNR), more reliable edges will be more similar across methods.

We computed the edge similarity as correlation at every edge for each pair of FC methods. We estimated the test-retest reliability using the intraclass coefficient (ICC) for each method separately. We estimated the variance components within the linear mixed model framework using the restricted maximum likelihood (REML) procedure [56, 57]. We defined variance components as follows:

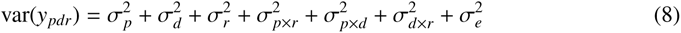

where *y* is an estimate of an edge, *p* indicates participant, *d* day, *r* run and *e* residual.

We computed the ICC as a ratio between between-subject variance (which included interaction terms pertaining to participants) and the total variance [58]. For this analysis, the runs were not concatenated.

Finally, we applied Fisher’s *z*-transformation to both edge similarity and ICC and computed the correlation between them. To reduce the number of comparisons, we only investigated the most relevant comparisons: full correlation vs. lagged correlation, partial correlation vs. multivariate AR1, and partial correlation vs. multivariate AR1 without self-connections. Since we estimated test-retest reliability separately for each method in a pair, there were two correlations for each pair of methods. We averaged both correlations for each comparison.

### 2.6. Node centrality measures

In the second part, we compared FC estimates using four different centrality measures: mean strength, eigenvector centrality, PageRank centrality, and participation coefficient. It is important to note that we did not use path-based methods, such as betweenness centrality or closeness centrality, because their interpretation is not clear for correlation-based networks. In such networks, a statistical association between two nodes does not necessarily indicate a path of information flow [1, 59]. Moreover, the correlation coefficient already captures the shortest path between two nodes [1].

The mean strength was computed as a mean of the edge weights for each region and it is analogous to a degree (number of connections) in binary networks. The shortcoming of strength is that it gives equal weight to all connections – it gives equal importance to nodes that are connected to other important nodes and to nodes that are connected to unimportant nodes.

In contrast, eigenvector centrality also considers the importance of a node’s neighbors. We computed the eigenvector centrality of a node *i* as the *i*-th entry of a principal eigenvector of the network’s adjacency matrix [60, 1]. Using a recursive formula, the eigenvector centrality can be expressed as:

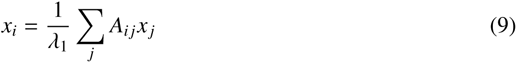

where, *x_i_*is the centrality of the *i*-th node, λ_1_ is the principal eigenvalue, and *A_i j_*are the elements of the adjacency matrix.

Eigenvector centrality has some drawbacks. For example, a node will be assigned zero centrality, if all of its neighbors have zero centrality. Additonally, a node with high centrality will give all of its neighbors a high centrality score, even if this is not intuitively meaningful. Consider a network of websites – if a website is indexed by Google, it will be assigned a high centrality, even if it has no other (incoming) connections. PageRank centrality was developed to address these limitations:

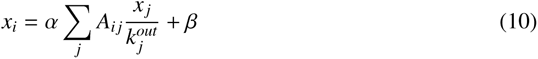

A positive constant *β* (usually set to 1) is added to ensure that no node has zero centrality and *x _j_* is divided by the out-degree of node *j* (*k_j_^out^*) to prevent high-degree nodes from having a disproportionate influence on other nodes [60, 1]. The balance between the eigenvector and the constant term is controlled by the parameter α, which is usually set to 0.85.

We computed both eigenvector and PageRank centrality using the implementations available in the Brain Connectivity Toolbox [59].

The degree-based or strength-based measures may be biased in correlation networks based on Pearson correlation, as node strength tends to correlate with community size [61, 62]. To mitigate this bias, we additionally used the participation coefficient to characterize node importance.

The participation coefficient measures the distribution of a node’s connections across different modules [63]. If the node’s connections are evenly distributed across modules, the participation coefficient approaches 1, while a participation coefficient of zero indicates that the node’s connections are completely restricted to its module.

The original formulation of the participation coefficient [63] does not take into account the size of the module [64, 62]. In particular, nodes from small modules tend to have high participation coefficients, while nodes from large modules tend to have low values. Therefore, we used the normalized participation coefficient [62]:

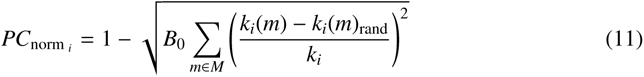

Here, *M* is a set of modules (communities), *k_i_*is the total degree of node *i*, *k_i_*(*m*) is the intramodular degree for node *i* in module *m*. *k_i_*(*m*)_rand_ represents a median intramodular degree for node *i*, obtained by generating randomized networks using the Maslov-Sneppen rewiring algorithm [65]. *B*_0_ is a multiplicative term used to constrain the range of *PC*_norm_ between 0 and 1 and was set to 0.5. The number of randomizations was set to 100 at the individual level and to 1000 at the group level. The module definitions were taken from Ji et al. [66].

We calculated centrality measures at the level of individual FC matrices and at the level of the group-averaged FC matrix. We then compared centrality measures based on different FC methods by computing Pearson’s correlation between the obtained centrality measures. For the comparison at the individual level, we averaged the obtained correlations. Additionally, to better understand the relationship between different centrality measures, we computed correlations between different centrality measures for selected FC methods (full correlation, partial correlation, and multivariate AR model).

In the case of dynamic FC estimates, all centrality measures were estimated separately for incoming and outgoing connections. The matrix of outgoing connections was obtained by transposing the original FC matrix. In addition, all centrality measures were estimated separately for positive and negative connections. For the sake of brevity, only the results for positive connections are presented in the main text; other results can be found in the Supplement. We refer to the strength of nodes based on incoming or outgoing connections as in-strength and out-strength, respectively.

### 2.7. Brain-behavior associations

To compare the brain-behavior associations obtained by different FC measures, we used 58 behavioral measures (see Table S1) that included cognitive, emotion and personality measures and were previously used in other studies [32, 67, 68].

#### 2.7.1. Variance component model

We computed brain-behavior associations using the multivariate variance component model (VCM), developed by Ge et al. [69] to estimate heritability. The use of the variance component model to estimate associations between the brain and behavior was introduced by Liégeois et al. [32]. We adopted the same approach to allow direct comparison with the results reported by Liégeois et al. [32]. Furthermore, the use of VCM allows an easy calculation of the explained variance for single traits. The model has the form

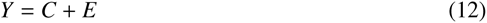

where *Y* represents the *N × P* matrix (number of subjects × number of traits) of behavioral measures, C represents shared effects and E represents unique effects. The model has the following assumptions:

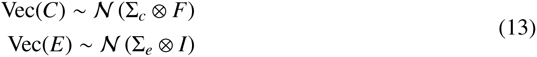

where Vec() is the matrix vectorization operator, is the Kronecker product operator, and *I* is the identity matrix. *F* represents *N × N* matrix of similarities between participants, which were estimated with the Pearson’s correlation coefficient. Σ*_c_*and Σ*_e_* are *P × P* matrices, which are being estimated. The total variance explained is computed as:

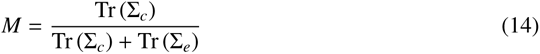

where Tr(·) represents the trace operator, and:

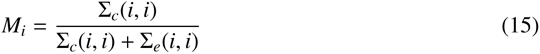

for single traits. *M* is analogous to the concept of heritability and can be interpreted as the amount of variance in behavior that can be explained with the variance in the connectome.

Before computing VCM, we imputed missing behavioral data using the R package missForest [70]. There were 0.59% missing data points overall. Following the procedure of Liégeois et al. [32], we applied quantile normalization to behavioral data. To remove potential confounding factors, we regressed age, gender, race, education, and movement (mean FD) using the procedure described in Ge et al. [71, 69].

We estimated *M* for each connectivity method separately. We compared patterns of explained variances by correlating the variance explained at the trait level between all methods.

Since the results of VCM are based on similarities between participants (matrix *F*), we tested the extent to which the similarities between participants, and thus the results of VCM, depend on the levels of noise in the data. To this end, we performed a simulation in which we added random Gaussian noise (mean 0, standard deviation 0–1 in steps of 0.1) to the standardized time series. To reduce complexity, we performed this analysis only for static FC methods.

#### 2.7.2. Canonical correlation analysis

Since VCM is rarely used to study brain-behavior associations, we repeated the analysis using canonical correlation (CCA). CCA is used to reveal the low-dimensional structure of the shared variability between two sets of variables (in our case, connectivity and behavior).

Let *X* and *Y* be *N × P* and *N × Q* matrices (*N* is the number of observations, *P* and *Q* are the number of variables), respectively.

The goal of CCA is to solve the following system of equations:

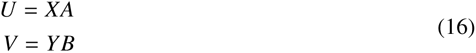

Here, *U_N_*_×*K*_ and *V_N_*_×*K*_ represent matrices of canonical scores (or variables), and *A_P_*_×*K*_ and *B_Q_*_×*K*_ represent matrices of canonical weights. The objective is to maximize the correlation between pairs of columns from *U* and *V* with the same index. These correlations are known as canonical correlations. The solution to the above set of equations is found under the constraint *U*^′^*U* = *V*^′^*V* = *I*. The columns of the *U* and *V* matrices tell us the relative position of each observation in the canonical variables. Columns of the *A* and *B* matrices contain information on the relative contribution of each variable to each of the canonical variables. Additionally, one can calculate canonical loadings - the correlations between original data matrices and canonical scores. Canonical variables are ordered in descending order according to the size of canonical correlations. Usually, only the first or first few canonical components are of interest, as these explain most of the shared variance. Mathematical details on CCA can be found elsewhere [e.g. 72, 73, 74, 75, 76].

We performed the CCA using the GEMMR package [73]. To prepare the data for CCA, we followed the procedure by Smith et al. [77], including deconfouding using the same variables as for VCM. Prior to CCA, we reduced the dimensionality of both sets of variables to 20 components using principal component analysis (PCA). This number was chosen to optimize the number of samples per feature based on the recommendation by Helmer et al. [73] under the assumption of a real first canonical correlation *r* = .30. We performed a 5-fold cross-validation to assess the generalizability of the model. We only examined the first canonical correlation since it was shown that the first canonical variable explains the most shared variance, and it was the only statistically significant canonical variable in a previous study [77].

We repeated the CCA for all FC methods. The similarities between the methods were assessed by comparing the first canonical correlation obtained in the training and the test set. Next, we correlated the canonical weights and loadings related to behavior.

#### 2.7.3. Principal least squares

Finally, we used principal least squares (PLS) to estimate brain-behavior associations. PLS is similar to CCA, with the goal to maximize covariance rather than correlation between sets [73, 78]. When the columns of *X* and *Y* are standardized, PLS gives the same results as CCA.

It has been shown that the first PLS component is biased toward the first principal component (PC) axis [73]. To assess the degree of bias in our data, we estimated the similarity between the PLS/CCA weights or loadings for behavior and the weights for the first behavioral PC.

#### 2.7.4. Control analyses

Participants in the HCP dataset are genetically and environmentally related, which can inflate between-subject similarities and influence the results related to interindividual differences. Therefore, we repeated all analyses related to brain-behavior associations on two subsamples of genetically unrelated participants (sample sizes 384 and 339).

### 2.8. Simulation

We hypothesized that dynamic and static FC estimates would be similar due to autocorrelation of fMRI time series, which is partly the result of convolution of neural time series with HRF. In addition, an important source of similarities (or differences) between FC results obtained by different methods could be due to similar (or different) effects of the amount of noise and the amount of available data on the resulting FC matrices. To evaluate the impact of convolution with HRF, signal quality, and the amount of data on estimated similarities between results using different FC measures, we used numerical simulations of data with known covariance structure. We generated multivariate time series of events for 1000 ”participants.” Events were sampled from a multivariate normal distribution with a mean of zero. The covariances differed for each participant and were taken from experimental data parcellated using Schaefer’s local-global parcellation with 100 regions [79]. We used this parcellation instead of MMP to reduce the computational burden and the size of the generated data. Events were not autocorrelated. The generated events were then convolved with HRF using the SimTB toolbox [80]. TR was set to 0.72 s (the same as in HCP data), and HRF parameters were set equal for all participants and regions (delay of response: 6, delay of the undershoot: 15, dispersion of the response: 1, dispersion of the undershoot: 1, the ratio of response to the undershoot: 3, onset in seconds: 0, length of the kernel in seconds: 32). The resulting time series were standardized.

To estimate the effects of signal quality on FC estimates and on similarities between FC methods, we added Gaussian noise with zero mean and standard deviation ranging from 0 to 1 standard deviation in steps of 0.1. This translates to SNR from 10 to 1 (excluding time series without noise, which has infinite SNR). We varied the time-series durations from 500 to 10000 data points in steps of 500.

The first step in the analysis was to establish the ground truth for each method, that is, the results that would be obtained in an ideal situation. We defined the ground truth as FC at maximum length and without noise in the event time series. Note that because events were not autocorrelated, the ground truth for all autoregressive FC methods was a matrix with all zero entries.

Next, we compared results using different FC methods in the same manner as for experimental data for all noise level and signal length combinations on prewhitened and non-prewhitened data. We computed (1) correlations between ground truth FC matrices and simulated FC matrices for all FC methods and (2) correlations between FC estimates obtained using different methods. To reduce the number of comparisons, we only investigated the most relevant comparisons: full correlation vs. lagged correlation, partial correlation vs. multivariate AR, and partial correlation vs. multivariate AR without self-connections.

## 3. Results

### 3.1. Similarities between FC estimates obtained using different methods

To address our research questions, we first focused on estimating similarities between the results obtained with different FC methods using empirical data. Comparison of group-level FC matrices showed very high correlations between FC results obtained using bivariate methods (*r* ⩾ .87, Figure 2A), as well as between results obtained using multivariate methods (correlation between partial correlation [AR1 prewhitened] and multivariate AR model: *r* = .80). In contrast, the correlations between the bivariate and multivariate FC estimates were lower and ranged from .36 to .65.

When comparing and pooling results based on individual-level FC matrices, the mean correlation between FC matrices obtained using different methods was lower. The correlations between the bivariate methods were still very high (correlation between lagged and full correlation: *r* = .99, correlation between prewhitened lagged and prewhitened full correlation: *r* = .83), while the correlations between the multivariate methods were lower on average. In particular, the correlation between the partial correlation (AR1 prewhitened) and the multivariate AR model was .05, compared to .80 between the group-level FC matrices.

The correlations between the results obtained using static and dynamic FC methods were smaller after prewhitening, with the greatest differences when comparing individual-level FC matrices obtained using multivariate methods. Specifically, the correlation between the coefficients of the multivariate AR model and the partial correlation decreased from .40 to .05 in the individual-level FC and from .86 to .80 in the group-level FC. The order of prewhitening had minimal effect on the correlations between the methods (Figure S2A), except for the comparison of the results obtained using the multivariate AR model and the partial correlation at the individual-level FC, where the correlations increased from .05 to .12 (*r* = .15–.22 for the multivariate AR model without self-connections).

The correlations between the FC results obtained using different methods were slightly higher when the analysis was repeated on 200 participants with the highest data quality (Figure S3).

#### 3.1.1. Autocorrelations of fMRI time series

To test the prediction that the similarities between the dynamic and static FC estimates would be similar to or greater than the mean autocorrelation of the fMRI time series, the mean autocorrelation function was computed across all participants and regions. The autocorrelation before prewhitening was .40 at lag 1 (Figure 2B). This autocorrelation decreased to .10 after AR1 prewhitening, and was essentially zero after AR2 and AR3 prewhitening. Thus, the similarities between the dynamic and static FC were always higher than the autocorrelation at lag 1. Interestingly, prewhitening at orders 1 and 2 reversed the sign of the autocorrelation at low lags.

### 3.1.2. Correlation between edge similarity and test-retest reliability

We computed edge similarity between FC methods as a correlation over subjects at every edge for selected pairs of FC methods. We estimated test-retest reliability at every edge for each method separately. Next, we computed the correlation between edge similarity and test-retest reliability for each of selected pairs of FC methods. The correlation was moderate to high for pairs of multivariate methods (*r* = .47–.66) and high for pairs of bivariate methods (*r* = .55–.79, Figure 3). Prewhitening lowered the correlations.

**Figure 3:**
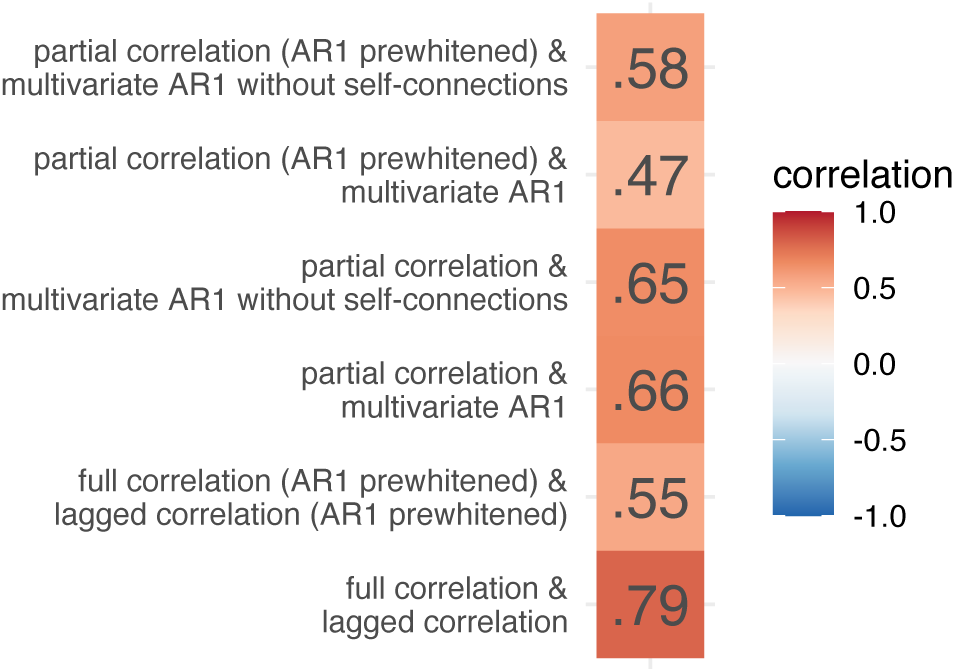
Correlations between edge similarity and test-retest reliability for selected pair of FC methods.

### 3.2. Similarity of node centrality measures

In the second part, we compared methods for estimating FC by comparing four node centrality measures: strength, eigenvector centrality, PageRank centrality, and participation coefficient (Figure 4, Figure S4, Figure S5). Unless otherwise noted, we focus on the positive connections. As before, we observed a clear distinction between bivariate and multivariate methods for computing FC matrices. Correlations between centrality measures based on bivariate FC methods were consistently high, regardless of the centrality measure (above .97 at the group level and above .96 at the individual level). In contrast, the correlations between multivariate FC methods were lower, ranging from .59 to .99 for strength, eigenvector centrality, and normalized participation coefficient at the group level. Correlations for PageRank centrality were generally lower for multivariate methods, ranging from .21 to 1.00.

**Figure 4:**
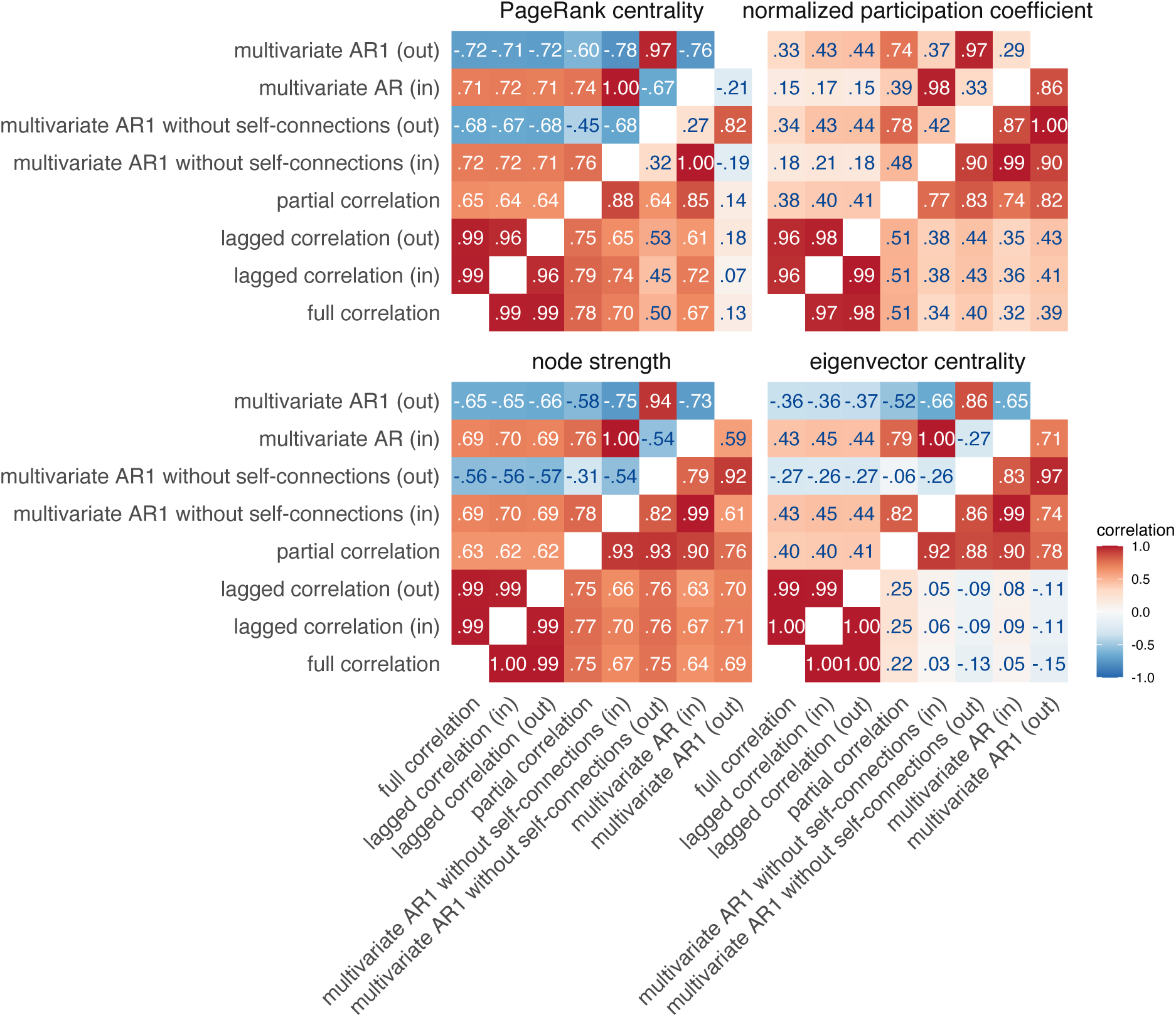
Similarities between node centrality measures based on positive connections. Similarities were estimated by (i) computing node measures on group-averaged connectivity matrices (group-level comparison; below diagonal), (ii) by computing node measures for each individual separately, correlating within participants and averaging these correlations across participants (individual-level comparison; above diagonal).

Similarities between multivariate and bivariate FC methods were moderately high for strength, PageRank centrality, and normalized participation coefficient, ranging from .32 to .79, except for PageRank centrality based on outgoing connections of the multivariate AR model, where the correlations were around .10. However, for eigenvector centrality, we found low similarities between multivariate and bivariate FC methods at the group level (ranging from .15 to .25) and moderate similarities at the individual level (ranging from .27 to .45).

Notably, we observed positive correlations between incoming and outgoing connections for strength-based centrality measures at the group level, but negative correlations at the individual level. This pattern was present only for multivariate dynamic methods. Additionally, when comparing partial correlation and multivariate AR models, we found that the correlations between strength-based centrality measures were positive for incoming connections and negative for outgoing connections. In contrast, all correlations were positive for the normalized participation coefficient.

We also found that our results were generally consistent when analyzing centralities computed on negative connections (Figure S5). More specifically, similarities between multivariate FC methods were smaller for normalized participation coefficient, and similarities between multivariate and bivariate methods were larger for eigenvector centrality.

Prewhitening reduced similarities between methods, especially for outgoing connections (Figure S4, Figure S5). This was evident for both positive and negative connections.

To better understand the similarities and differences between the FC methods, we plotted the distributions of the centrality measures on the cortical surface (Figure 5, Figure S8). For both static FC methods, the strength was highest in the parietal regions (Figure 5A). For partial correlation, eigenvector centrality was distributed similarly to strength, whereas for full correlation, the highest eigenvector centrality values were in visual and somatomotor cortex (Figure 5B). For partial correlation, participation coefficient values were lowest in visual and somatomotor cortex, and highest in frontal and parietal regions belonging to parts of the cingulo-opercular, dorsal attention and multimodal networks (Figure 5C). For the full correlation, similar to the partial correlation, the participation coefficient values were lowest in medial frontal regions and parts of visual cortex, and highest in parts of parietal cortex.

**Figure 5:**
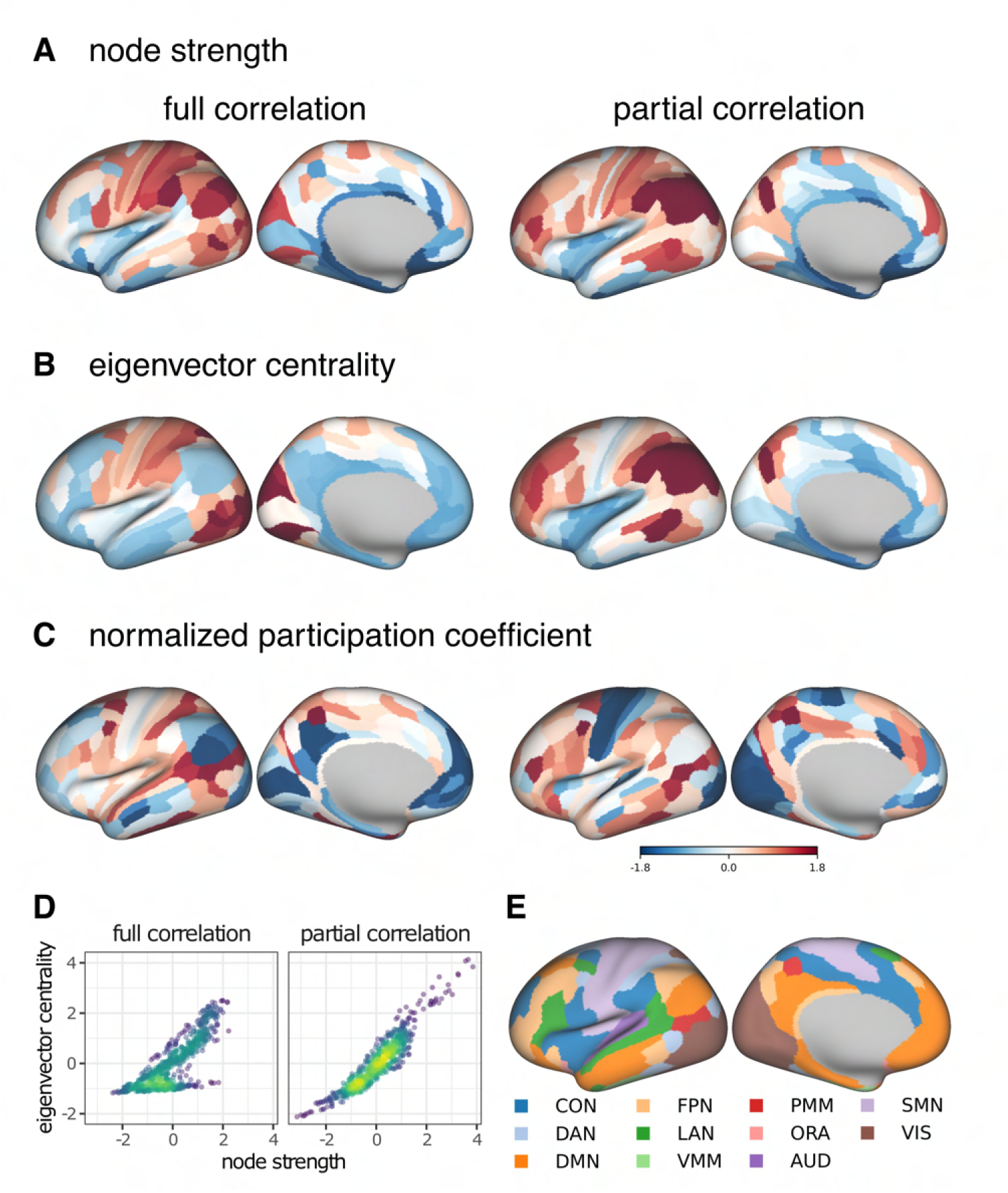
Cortical distribution of centrality measures based on static FC methods for positive connections. PageRank centrality is omitted, because its correlation with strength is close to 1. For visualization, the values have been transformed into *z*-values. **D.** Correlation between node strength and eigenvector centrality. **E.** Functional networks as defined in Ji et al. [66]. CON – cingulo-opercular network, DAN – dorsal attention network, DMN – default mode network, FPN – frontoparietal network, LAN – language network, VMM – ventral multimodal network, PMM – multimodal network, ORA – orbito-affective network, AUD – auditory network, SMN – somatomotor network, VIS – visual network.

We also plotted the distributions of centrality measures based on incoming and outgoing connections of a multivariate AR model (Figure 6, Figure S9). In-strength was highest in the parietal lobe, while out-strength was highest in the parts of the temporal lobe and in the medial parietal lobe (Figure 6A). Eigenvector centrality was similarly distributed for incoming and outgoing connections, with highest values in the frontal and parietal regions, mainly in parts of the frontoparietal network (Figure 6B). Similar to partial correlation, participation coefficient values were lowest in the somatomotor and visual cortex and highest in medial frontal and medial temporal cortex (Figure 6C). For incoming connections, the PageRank centrality was distributed similarly to the eigenvector centrality, while for outgoing connections, the values were highest in the medial parietal and medial temporal lobes, and lowest in the somatomotor and frontal regions (Figure 6D).

**Figure 6:**
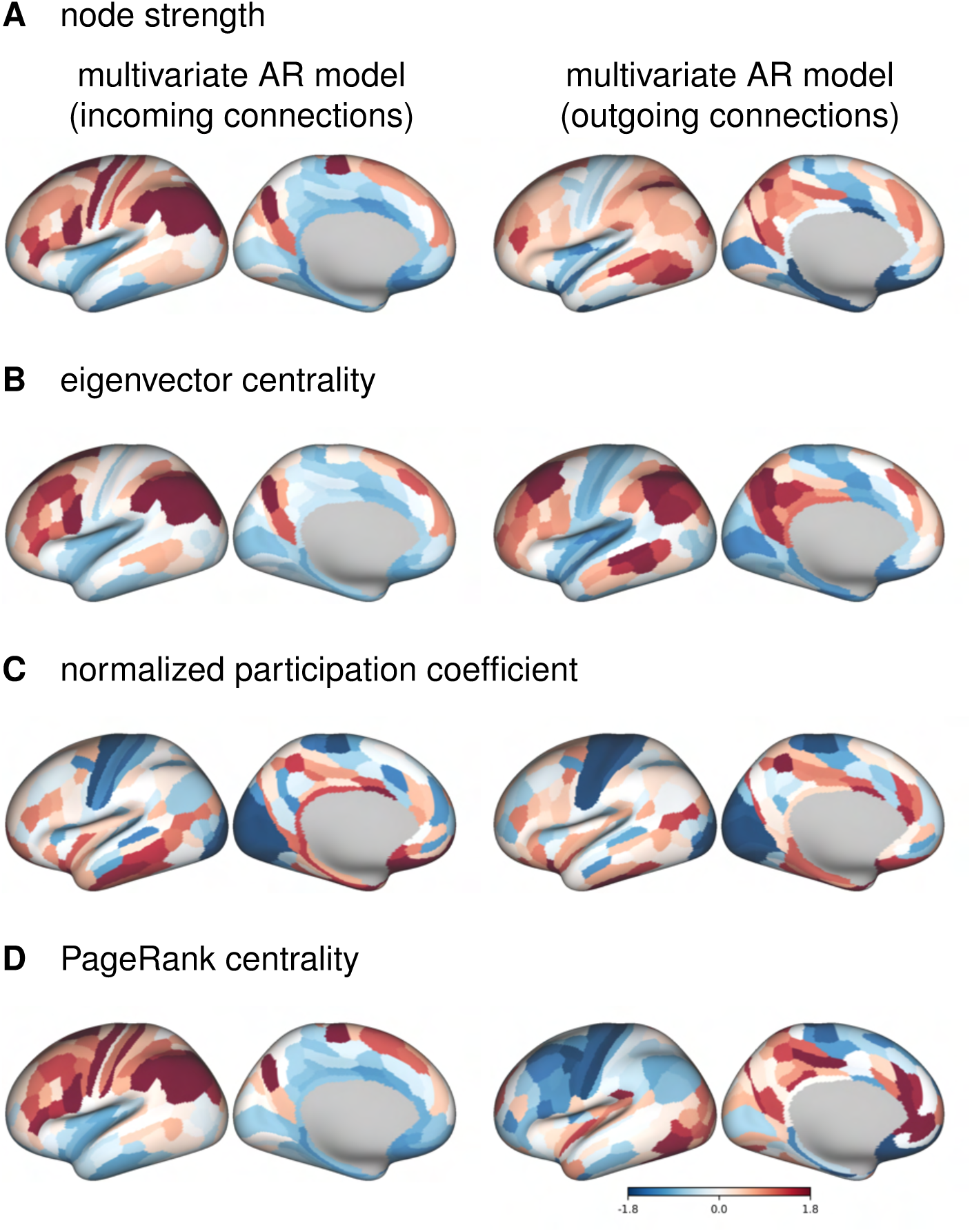
Cortical distribution of centrality measures based on multivariate autoregressive model for positive connections. For visualization, the values have been transformed into *z*-values.

Since in the case of the multivariate AR model the results of group-averaged connectomes differ from individual connectomes, we also plotted the distributions of the centrality measures for a representative subject (Figure S10, Figure S11). In the individual case, the distribution of strength-based centralities for outgoing connections is the opposite of that for incoming connections, with the lowest values in the parietal and occipital cortex.

Next, we analyzed the correlations between the centrality measures for full and partial correlation (Figure 5D, Figure S6). The correlations between the strength-based centrality measures were generally very high (*r* ⩾ .90) within positive and within negative connections. Exceptions were the correlation between eigenvector centrality and strength (*r* = .80) and the correlation between eigenvector centrality and PageRank centrality (*r* = .64) for positive connections on connectomes based on full correlation. In both cases, examination of the scatter plots (Figure 5D) revealed two groups of nodes – one group of nodes had higher similarity between centrality measures, while the other had a lower similarity. This pattern was also observed for negative connections, but with a smaller difference between the two groups.

### 3.3. Brain-behavior associations

Next, we compared patterns of brain-behavior associations derived from different FC methods.

#### 3.3.1. Variance component model

The results of the VCM show that bivariate methods explain about 30 percentage points less variance in behavior than multivariate methods (Figure 7A,B). Furthermore, the similarity of patterns of variance explained over behavioral measures was high between static and dynamic FC methods using the same number of variables, i.e., between full correlation and lagged correlation (*r* = 1.00), and between partial correlation and multivariate AR models (*r* = .83–.86, Figure 7A,C). The pattern of similarities in behavioral variance explained between the FC methods was comparable to the direct comparison of the FC matrices (Figure 7C, cf. Figure 2A). Patterns of similarities between the FC methods were similar when the analysis was performed on subsamples of unrelated participants (Figure S12A,C); however, the differences in total variance explained between the bivariate and multivariate methods were smaller (Figure S12B).

**Figure 7:**
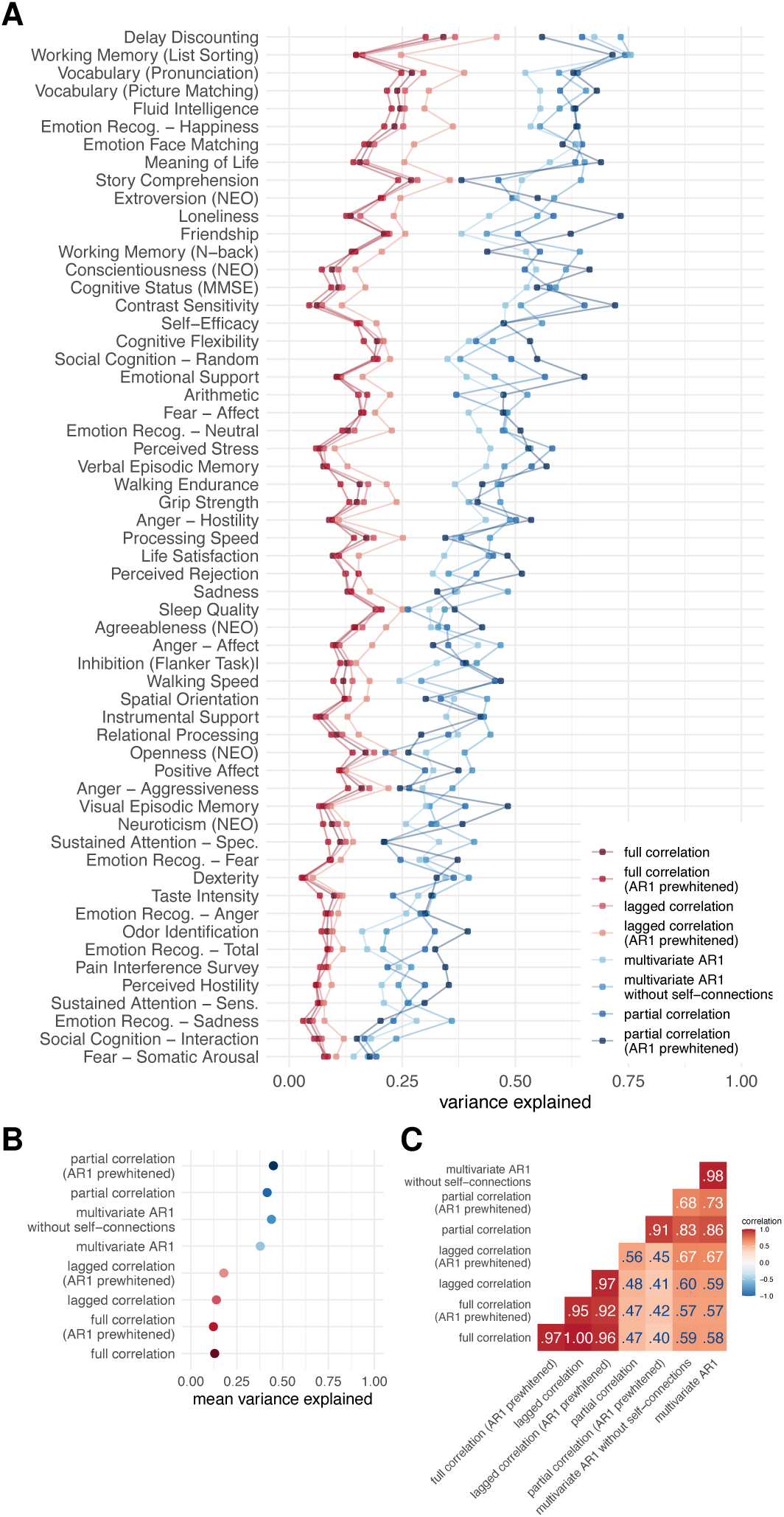
Results of variance component model for brain-behavior associations. **A.** Variance explained for individual traits estimated with different connectivity methods – traits are ordered according to the mean variance explained across connectivity methods. **B.** Mean variance explained. **C.** Similarities of explained variance patterns between connectivity methods.

Simulation of the effects of noise in which we added various levels of noise to the fMRI time series showed that noise affects estimates of the behavioral variance explained by the connectome. In particular, the mean of the variance explained increased with increasing noise for both the full correlation and the partial correlation, but the increase was more pronounced in the case of partial correlations (Figure 8B). This pattern was not equal for all behavioral variables – for some, the variance explained decreased and for others, it increased (Figure 8A). On the other hand, the similarity between the participants decreased with increasing noise (Figure 8C). This effect was more pronounced for partial correlation compared to full correlation.

**Figure 8:**
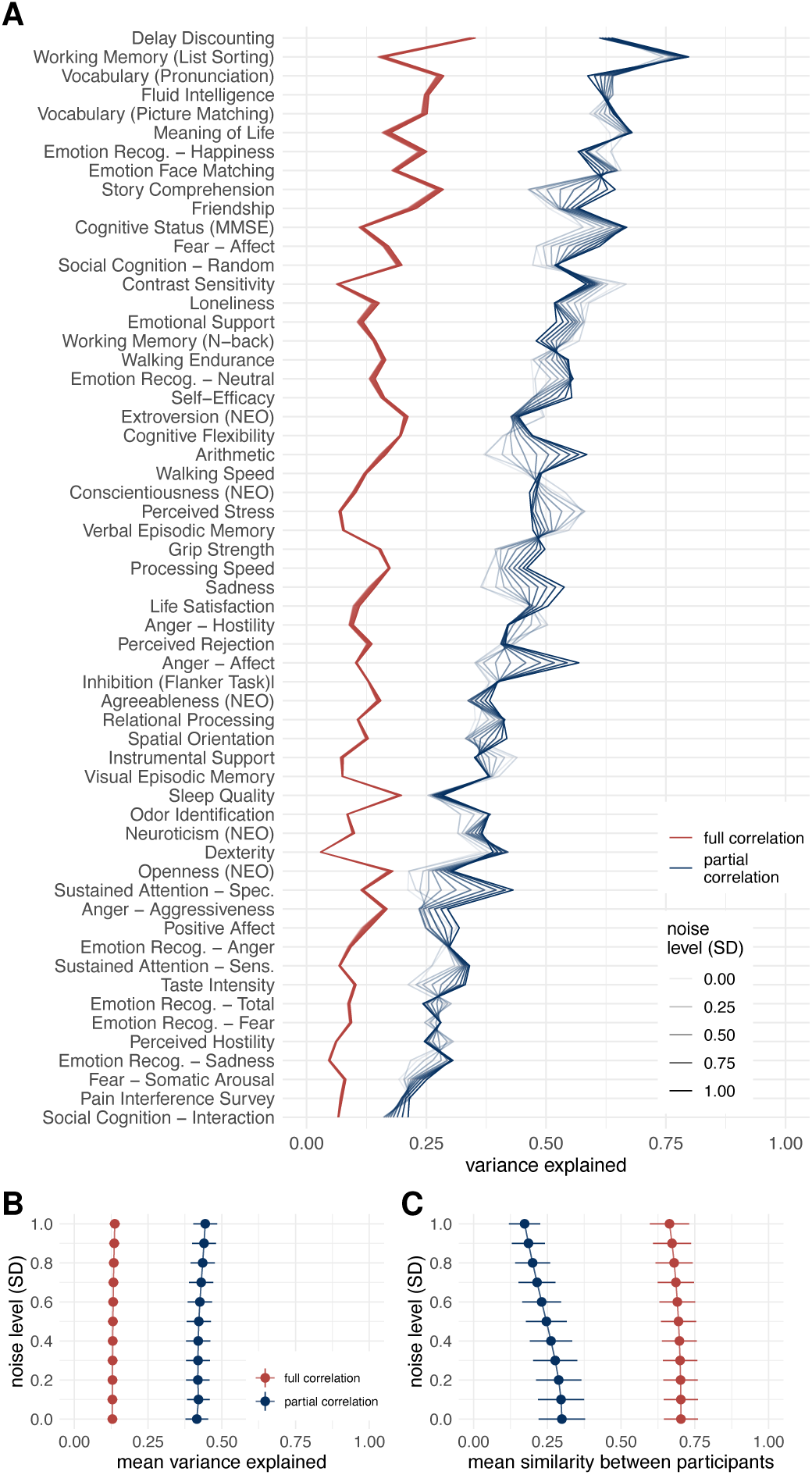
Results of variance component model for brain-behavior associations on data with added noise. FC was estimated using Pearsons’/full correlation and partial correlation after adding various levels of random Gaussian noise to experimental time series. **A.** Variance explained for individual traits estimated with different connectivity methods. Traits are ordered according to the mean variance explained across connectivity methods. **B.** Mean variance explained. Error bars represent jackknife standard deviation. **C.** Mean similarity between participants. Error bars represent standard deviation.

#### 3.3.2. Canonical correlation analysis

The results of the similarity between the FC methods when investigating brain-behavior associations using CCA were comparable to those obtained using VCM. In particular, the correlations between the weights or loadings on behavioral measures between the FC methods were high when comparing the methods that use the same number of variables for the estimation of a single edge (*r* > .80) (Figure 9C). On the other hand, there was no discernible difference between dynamic and static FC estimates.

**Figure 9:**
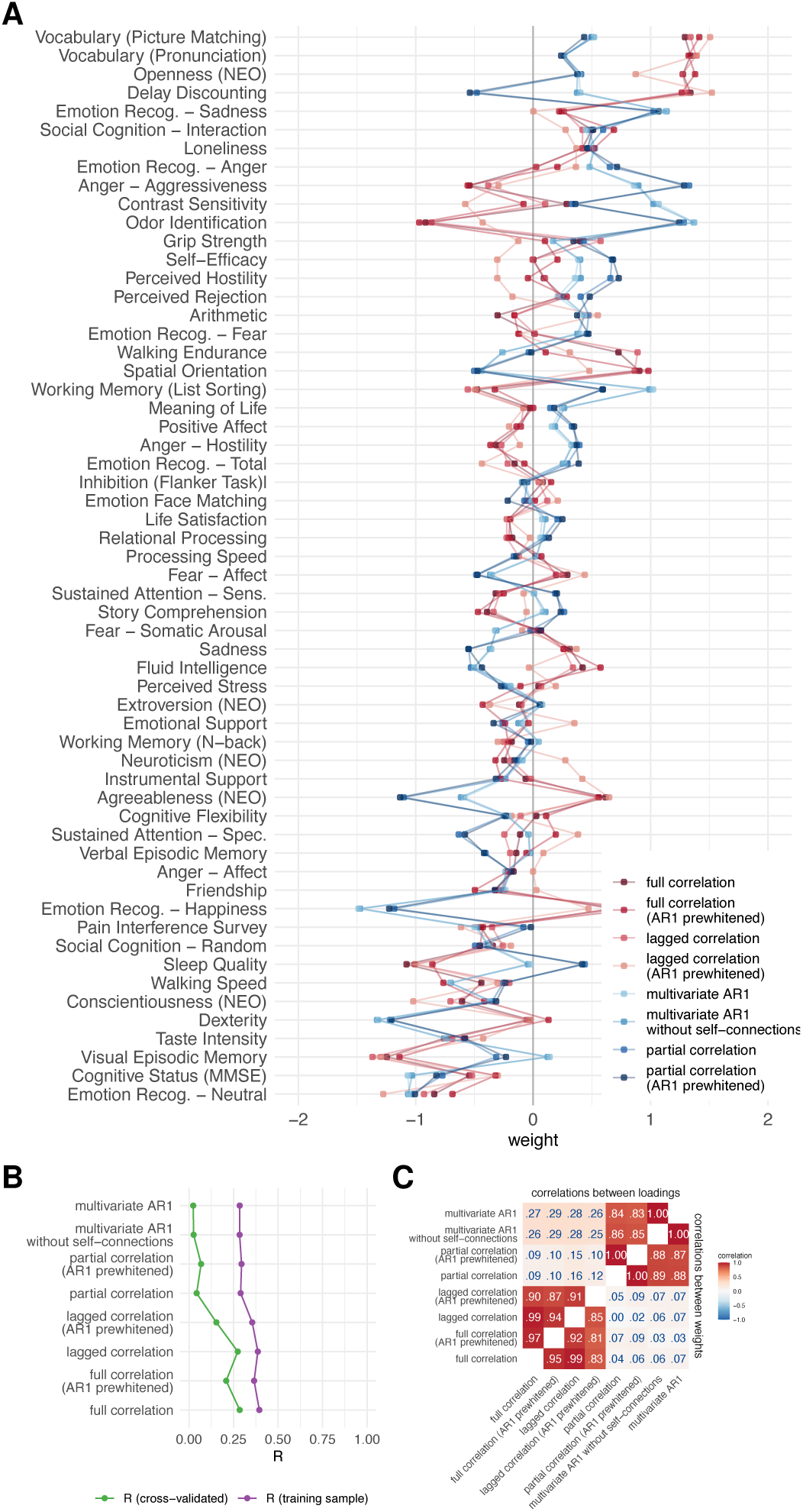
Results of canonical correlation analysis for brain-behavior associations. **A.** CCA weights. **B.** First canonical correlation on test and training set, **C.** Correlations between canonical loadings and weights across functional connectivity methods for first canonical components.

The first canonical correlation was around .35 in the training sample for the bivariate methods and around .30 for the multivariate methods (Figure 9B). Cross-validated R was much lower, around .25 for bivariate methods and around 0.05 for multivariate methods. Although these results differ from VCM (where multivariate methods explained more variance), the pattern of similarity between FC methods is the same.

The pattern of results was similar for the subsamples of unrelated participants (Figure S13B,D), but the differences between the training and test sets were larger (Figure S13A,C). The large difference between the performance of the model in training and test sets is indicative of overfitting, which is characteristic of CCA with a small number of samples per feature [73].

#### 3.3.3. Principal least squares

Analysis of brain-behavior associations using PLS revealed higher similarities between FC methods compared to CCA (Figure S14A,C). Specifically, the correlations between loadings were consistently greater than .91 for all methods compared, and the correlations between weights were greater than .51. Consistent with all previous results, we observed a clear separation between multivariate and bivariate methods when comparing weights, and no difference between static and dynamic FC methods based on the same number of variables. PLS was less generalizable compared to CCA, with canonical correlations on the training sample around .15–.20 and canonical correlations on the test sample around 0–.05.

However, our results also suggest that the high similarities between FC methods in PLS may be due to the strong similarity between the first behavioral canonical component and the first behavioral principal component, as reported in a previous study [73] (Figure 10). The correlations between loadings and the first behavioral principal component were around 1.00, while the correlations between weights and the first behavioral principal component were around .80. In contrast, for CCA, these correlations were about .60–.75 and .10–.25, respectively.

**Figure 10:**
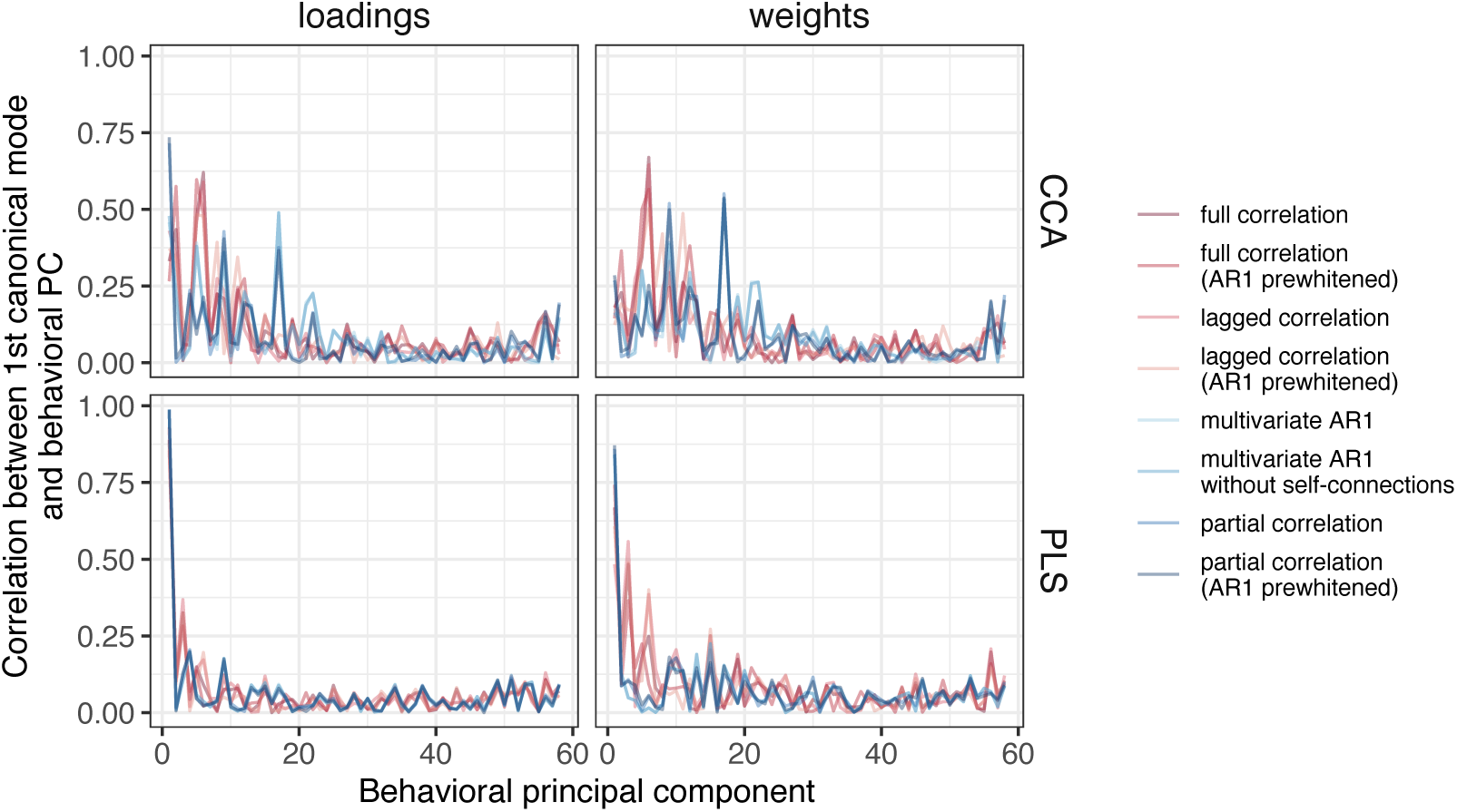
Similarity between first behavioral principal component and loadings or weights, for both CCA and PLS. For PLS, the correlation between loadings or weights and the first PC was very high. For CCA, the correlation with the first PC was high for loadings but moderate for weights. The differences between the FC methods were generally small, but somewhat larger for CCA than for PLS.

### 3.4. Evaluation of similarities between methods on simulated data

#### 3.4.1. Relationship between FC estimates and ground truth

Correlations of FC estimates with ground truth were greater than 0.8 for full correlation and between 0.25 and 0.9 for partial correlation (Figure 11B). Prewhitening decreased the correlation with ground truth. This effect was more pronounced for partial correlations. Longer time series also had higher correlations with ground truth (the difference was up to .5 for partial correlation and up to .3 for full correlation). The correlation with ground truth generally decreased with decreasing SNR (increasing noise), but in the case of partial correlation, these effects were not monotonic. In particular, for short time series, correlation with ground truth increased with low to moderate noise. Also in the case of partial correlation, prewhitening increased the correlation with ground truth at low noise. In contrast, prewhitening decreased the correlation with ground truth in the presence of high noise compared to the case without prewhitening.

**Figure 11:**
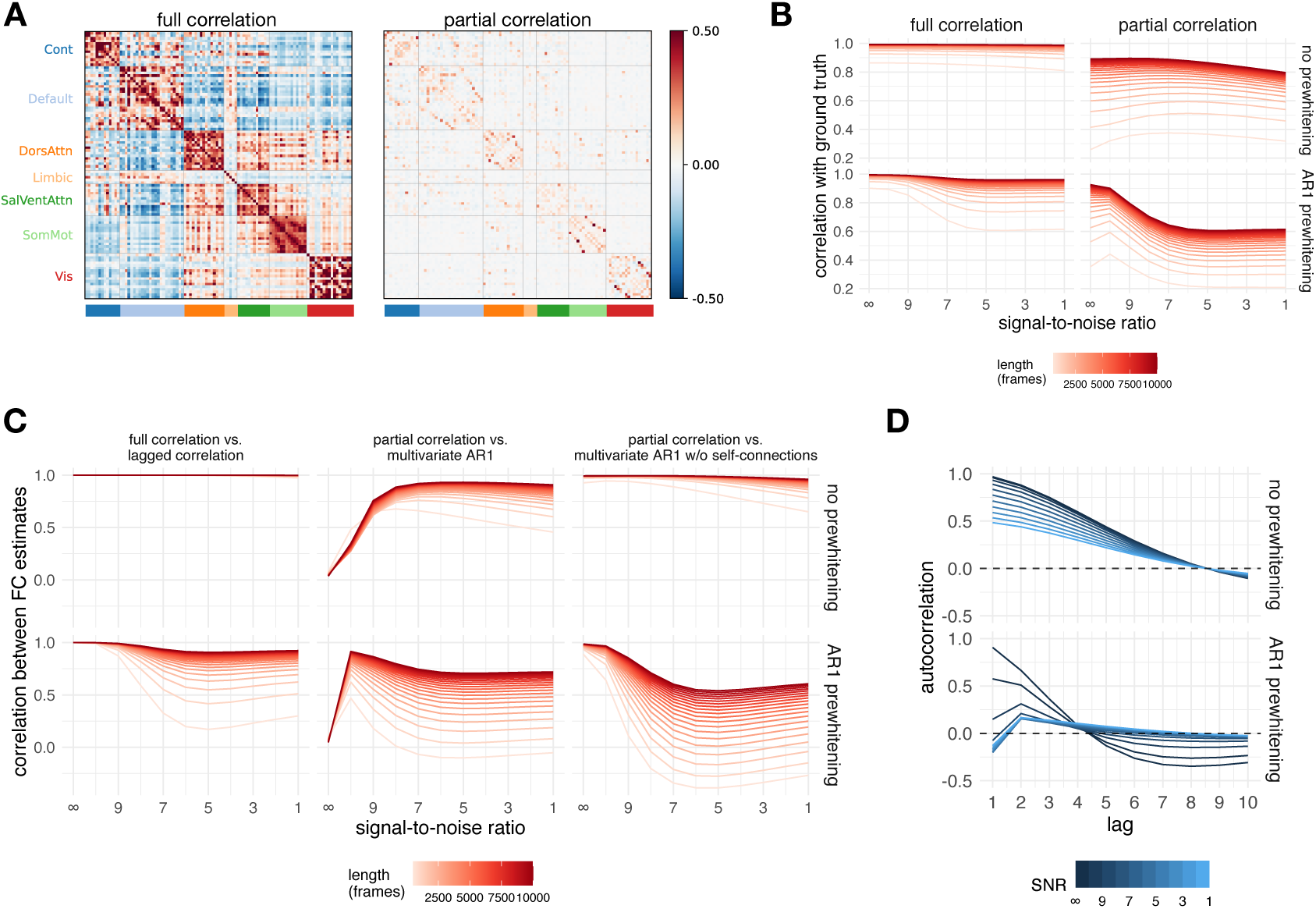
Results of simulation. **A.** Ground truth matrices (mean over participants). Note that all ground truth autoregressive model coefficients equal zero, since the simulated events were not autocorrelated. **B.** Correlation between the ground truth and the simulated data for all FC methods and their relationship to the noise level and signal length. **C.** Correlations between selected pairs of FC methods as a function of noise and signal length for simulated data. **D.** The autocorrelation function of the simulated data as a function of prewhitening order and noise.

#### 3.4.2. Similarity between FC estimates

The connectivity matrices computed on the simulated data were compared in the same manner as for the experimental data. For brevity, we focus only on the three most relevant comparisons (lagged correlation vs. full correlation, multivariate AR model vs. partial correlation, multivariate AR model without self-connections vs. partial correlation).

Estimates based on lagged and full correlation were highly similar (*r ≈* 1 in the case without prewhitening) for all levels of noise and signal length (Figure 11C). The correlation between FC estimates was reduced for prewhitened data, especially for low signal lengths (< 1000 frames).

The FC estimates of the multivariate AR model did not correlate with the FC estimates based on partial correlation when the noise was low (*r* = 0 for zero noise). However, with increasing noise and increasing signal length, FC estimates became very similar (up to *r* = .95), especially in the case without prewhitening and for long signal lengths.

Conversely, FC estimates based on a multivariate AR model without self-connections showed a high similarity to the FC estimates based on partial correlation at a low noise level (*r* > .95). For prewhitened data, there was a nonmonotonic relationship between FC estimates with increasing noise, but overall correlations remained high in conditions with high signal length.

For both multivariate AR models, the similarities to the partial correlation were negative for very short time series. This effect was more pronounced for higher levels of noise, but the relationship with noise was not monotonic.

To better understand the relationship between the multivariate FC methods we plotted the distribution of edge values as a function of noise separately for diagonal and off-diagonal terms (Figure S18). For brevity we did this only for the longest signal (10000 frames). For the multivariate AR model, the diagonal terms (self-connections) were close to 1 at very high SNR and decreased with decreasing SNR. Conversely, the mean of the off-diagonal terms remained close to zero, regardless of the SNR, but the variability increased with increasing SNR. The opposite pattern was observed when the data were prewhitened. At maximum SNR (i.e., when no noise was added to the data), the diagonal terms were essentially equal to one and the off-diagonal terms were essentially zero, with very low variability compared to all other distributions. The distribution of values at maximum SNR was not affected by prewhitening. For the multivariate AR1 model without self-connections and partial correlations, the variability of the edges decreased with increasing SNR. Prewhitening reduced the variability of the edges.

#### 3.4.3. Autocorrelation on the simulated data

We computed the average autocorrelation function over all participants and regions (Figure 11D, Figure S1). In general, noise and prewhitening reduced the absolute autocorrelation. The shape of the autocorrelation function varied as a function of noise and prewhitening. In the case without prewhitening, the autocorrelation decreased monotonically, reaching 0 at lag 8. With AR1 prewhitening, the autocorrelation decreased to negative values after lag 4.

## 4. Discussion

In this study, we addressed the question of whether the temporal order of the BOLD fMRI time series contains information important for the study of the fMRI brain functional connectivity. To this end, we compared FC estimates between methods that differ in their sensitivity to temporal order, i.e., static and dynamic measures of FC. We also compared methods that differed in the number of variables considered in estimating the connectivity of individual edges, i.e., bivariate and multivariate. Our results suggest that dynamic FC connectivity methods provide similar connectivity estimates as static FC methods of the same type (bivariate or multivariate), whereas bivariate and multivariate methods differ in terms of the explanation of individual differences in behavior.

### 4.1. Dynamic functional connectomes represent information similar to static functional connectomes

By directly comparing the FC matrices, we have shown that the estimates of the dynamic FC represent information similar to the estimates of the static FC. The similarity between estimates of FC, obtained by different methods, depended on several factors. First, there were high correlations between the FC estimates when the same number of variables was considered (Figure 2A). Second, similarities between connectomes were greater when averages were compared at the group level than when correlations were aggregated across individual-level FC matrices. We believe that the differences between the group- and individual-level cases are mainly due to better SNR in the case of the group-level data. Two observations support this conclusion: first, similarities in FC estimates between methods were greater for participants with the highest data quality, and this effect was more pronounced when comparing individual-level matrices than at the group level. Second, edges with higher test-retest reliability (an indicator of SNR) were more similar between FC estimates obtained by different methods. Thus, we can conclude that SNR influences the similarity between FC estimates.

Using simulation, we tested the similarities between FC as a function of noise and signal length (Figure 11C). We have shown that the dynamic FC estimates resemble static FC estimates even in the absence of true lagged correlation. The similarity between the multivariate AR1 model and partial correlations can be partially explained by the fact that the multivariate AR1 coefficients are a product of the inverse covariance and the lagged covariance matrix. In the case of the multivariate AR model, the similarity to the partial correlation was actually higher when more noise was added to the data. This occurs because the self-connections (the diagonal term in the AR matrix) act as a prewhitening term. When the SNR was maximal, the self-connections were close to 1 and the off-diagonal terms were close to zero (Figure S18). In other words, the self-connections explained all the variance in the time series and there was no variability left to be explained by the off-diagonal terms. When noise was added to the data, the autocorrelations were reduced (Figure 11D) and the self-connections shrank (Figure S18). Consequently there was less prewhitening due to the self-connections and the off-diagonal elements became more similar to the partial correlations. For the same reason, estimates based on a multivariate AR1 model without self-connections were highly correlated with estimates based on partial correlation regardless of the noise level – there were no self-connections to explain the autocorrelation.

We also found a high similarity between the full and the lagged correlation. Therefore, the similarity between the multivariate AR1 model and the partial correlation cannot be explained solely by the inclusion of the precision matrix in the estimation of the coefficients of the multivariate AR model. Rather, the lagged covariance matrix also contributes to this effect.

We hypothesized that the similarities between the dynamic and static FC estimates originate from autocorrelation of the fMRI time series. We predicted that the similarities between the dynamic and static FC estimates would be at least as large as the average autocorrelation of the fMRI time series and that this similarity would be reduced after prewhitening. Both predictions were confirmed in experimental and simulated data. However, even when autocorrelation was reduced to virtually zero at all lags (this occurred at prewhitening order 3), similarities between estimates based on dynamic and static FC models remained high for group-level matrices and simulated data. This suggests that prewhitening (or even the presence of noise that reduces autocorrelation) does not completely eliminate the influence of convolution with HRF on the estimation of dynamic FC.

We conclude that even if AR models represent information that goes beyond the static FC, this cannot be claimed on the basis of a direct comparison of dynamic and static FC estimates.

One of the main differences between static and dynamic FC methods is the ability of dynamic FC methods to estimate the directionality of connections [23]. FC matrices based on dynamic FC methods are therefore asymmetric. To allow comparisons between static and dynamic FC matrices, the former were symmetrized and the information about the directionality of the connections was lost. To test the possibility that there is specific information in the dynamic FC estimates that could not be detected in a direct comparison of the FC matrices, we additionally compared the node centrality measures and the patterns of brain-behavior associations between the FC methods.

### 4.2. Network topology is affected by the functional connectivity estimation method

Examining node centrality measures allowed us to investigate how different FC methods affect network topology. First, we analyzed the similarity between FC estimates based on different FC methods for each centrality measure separately. Overall, the results were consistent with direct comparisons of FC matrices. We found a clear distinction between multivariate and bivariate FC methods, while the difference between static and dynamic FC estimates was rather small (with an important caveat regarding the difference between incoming and outgoing connections, see below). The similarities were also influenced by the choice of the node centrality measure. In particular, the similarities between multivariate and bivariate FC methods were relatively low for eigenvector centrality (from .15 to .25 for the group-level comparison), while for other centrality measures the similarity between multivariate and bivariate methods was higher (e.g. around .70 for strength).

We explored this finding further by examining the similarities between the centrality measures. While the correlations between the centrality measures were predominantly positive, and in some cases close to 1, there were some exceptions, suggesting that the centrality measures are not redundant. Specifically, for the correlation between eigenvector centrality and strength computed on full correlation connectomes, we observed two groups of nodes, one with higher similarity between the two centrality measures and one with lower similarity (Figure 5D). This pattern has been observed before [81] and suggests that one group of nodes is connected to other important nodes, while the other is mainly connected to less connected nodes. In other words, these two groups of nodes can be distinguished by jointly considering both eigenvector centrality, which measures how well a node is connected to other important nodes (i.e., nodes with many or strong connections), and strength, which is affected only by the number or strength of a node’s connections. Notably, however, we observed this pattern for full correlation, but not for partial correlation. This suggests that indirect connections have an important influence on the global position of nodes in functional connectomes estimated using full correlation. When indirect connections are removed (i.e., when partial correlation is used to estimate FC), the topological position (importance) of a node is the same regardless of the centrality measure. In summary, the choice of FC method has a different impact on the network topology depending on the centrality measure used.

Second, we were interested in the relationship between incoming and outgoing connections. For multivariate AR model estimates, we found a negative correlation between in-strength and out-strength when comparing at the individual level. However, when comparing group-averaged FC matrices, the correlations between in-strength and out-strength were positive. Interestingly, when comparing the partial correlation with the multivariate AR model, the correlations of strength from the partial correlation connectomes were positive with in-strength from the multivariate AR model, but negative with out-strength. The individual-level results confirm previous findings [30], suggesting that brain regions are either feeders or receivers, but not both. However, this information is lost when FC matrices are averaged across subjects. In addition, there was a positive correlation between the in-strength and out-strength of bivariate dynamic FC estimates, regardless of the level of comparison. In FC analyses, individual-level matrices are often averaged, concatenated [e.g. 34], or estimated using a group prior [e.g. 82]. Because group-level FC matrices may be qualitatively different from individual-level FC matrices, we recommend that researchers perform analyses and/or examine results at both the group and individual levels whenever possible and/or meaningful.

### 4.3. Dynamic FC models do not explain additional variance in behavior over static FC models

We used the variance component model (VCM), canonical correlation analysis (CCA), and principal least squares (PLS) to estimate brain-behavior associations. The results of all methods showed that there were no large differences between the dynamic and static FC estimates in the patterns of associations with behavior. However, we found large differences between the bivariate and multivariate methods. These differences were specific to the method used to estimate brain-behavior associations.

In the case of CCA, the canonical correlations were higher for bivariate methods than for multivariate methods. The cross-validated canonical correlations for multivariate methods were around 0, indicating that the results were not generalizable. In contrast, the difference between the canonical correlations in the training and test sets was relatively small for the bivariate methods.

In the case of PLS, the similarities between the FC methods were extremely high, especially when we compared loadings. We showed that these results are most likely due to the high similarity of the behavioral loadings and weights to the first behavioral PC, confirming previous observations that the PLS loadings and weights are biased toward the first principal axis, especially at low sample-to-feature ratios [73]. Compared to PLS, CCA, on the other hand, shows much less bias toward the first principal axis. In addition, the canonical correlations based on PLS had negligible generalizability. Therefore, we advise users to be cautious when using PLS. We recommend that users perform cross-validation and examine the similarity between canonical weights/loadings and PCs.

In the case of VCM, the multivariate methods explained on average about 30 percentage points more variance in behavior than the bivariate methods. To better understand this observation, we examined the impact of inter-subject similarities on VCM results. To this end, we added random noise to the data, reducing the similarities between subjects. Interestingly, full correlation and partial correlation explained more variance in behavior on average when we added random noise to the data. This may sound counterintuitive, but keep in mind that VCM was developed to estimate heritability [69], that is, the proportion of variance in phenotype that can be explained by variance in genotype. Holding the environment constant, higher genetic similarity would reduce the estimate of heritability. If all individuals within a sample had the same genotype, heritability would be zero because no variance in phenotype could be explained by variance in genotype. The input to VCM is a between-subject similarity matrix (usually a genetic similarity matrix or, in our case, a connectome similarity matrix). Participants were more similar when we used full correlation as an estimate of FC compared to partial correlation. This explains the observation that the partial correlation explained more variance in behavior.

Our second simulation showed that the partial correlation estimates are less stable and more affected by noise and signal length. This explains the apparent discrepancy between VCM and CCA. Our results show that when we add noise to the experimental data, participants become more dissimilar and, in the case of VCM, the proportion of behavioral variance explained by the variance in the connectome becomes larger. In the case of CCA, lower SNR leads to lower and less generalizable canonical correlations for multivariate FC methods. For this reason, we recommend that great care be taken when estimating brain-behavior associations with measures that are sensitive to noise.

Liégeois et al. [32] have used VCM to compare brain-behavior associations between correlation and the multivariate AR model. They concluded that the dynamic FC explained variance in behavior beyond that explained by static FC. We have shown that these results are confounded by the mixing of two orthogonal properties of the FC methods: sensitivity to the temporal order of time points and the number of regions used to estimate a single edge. The difference between the explanatory value of the multivariate AR model and the full correlation is better explained by the difference between the multivariate and bivariate nature of the method than by the sensitivity to the temporal order of the time points.

### 4.4. Relationship between static, dynamic and time-varying functional connectivity

As explained in the Introduction, dynamic and time-varying FC encode different aspects of temporal information in FC. Based on previous research investigating resting-state fMRI, which showed that FC is largely stationary [7] and independent of cognitive content [83], we assumed stationarity of FC time series, and chose models of stationary static and dynamic FC as the basis of our study (as opposed to models of TVFC). Nevertheless, stationary FC is not incompatible with the presence of meaningful FC fluctuations [7, 21].

Brain states can be estimated using TVFC estimation methods such as hidden Markov models (HMM) or clustering of sliding window correlation (SWC) matrices [10, 9]. Brain states derived in this way have been studied in the context of tracking ongoing cognition and behavior, and also for predicting trait aspects of behavior, such as personality, psychopathology, and performance on cognitive tests [see reviews in 10, 9, 84]. Commonly used metrics derived from brain states include transition matrix (a matrix that encodes the probabilities of transitioning from one state to another), fractional occupancy (proportion of time spent in each state), and switching rate (the frequency of switching between states) [85, 86, 87]. In addition, some studies have quantified TVFC using edge variability metrics, such as edge variance or standard deviation [88, 89], amplitude of low frequency fluctuations (ALFF) [90], and excursions from a median time-varying correlation [19]. Edge variability has been shown repeatedly to be negatively correlated with the static FC [19, 90, 88, 89], suggesting that stronger edges have lower variability, and that variability of FC is partially redundant with the edge strength derived from static and stationary FC.

Several studies have compared TVFC with static and stationary FC in terms of behavioral prediction, showing that TVFC-derived metrics have differential or better predictive power over static/stationary FC and/or anatomical brain features [87, 91, 31, 92, 86], and also for the prediction of psychopathology. Jin et al. [31] compared the predictive value of static, dynamic, and time-varying FC, and showed that TVFC had the best predictive value for PTSD. However, consistent with our findings, dynamic FC was only slightly better than static FC. Note that some of these studies suffer from methodological shortcomings, such as small sample sizes [87, 91], and thus the results may have low generalizability [73, 93]. Nevertheless, overall, the results suggest that TVFC does contain additional information beyond static or dynamic FC. Further studies are needed to reconcile these findings with the evidence that resting-state connectivity is largely stationary.

### 4.5. Limitations and future directions

A number of limitations should be considered in drawing conclusions from our study. First, in our simulation, we generated noise using a multivariate normal distribution. We could have used more advanced noise modeling that incorporated specific noise components such as drift, moving average, physiological noise, and system noise [94]. Unlike white noise, these noise sources are autocorrelated and therefore could affect the (dynamic) FC estimates. We wanted to keep the model simple and interpretable. Even with the simplest noise model without autocorrelation in neural time series, we showed that AR models can be affected by convolution of the neural signal with HRF and that consequently the dynamic FC estimates resemble the static FC. However, more advanced noise modeling could be used for a more realistic assessment of the sources of similarities between different FC methods.

Similarly, we used a very simple procedure, prewhitening, to reduce autocorrelation. Other methods could also be used to reduce autocorrelation, such as advanced physiological modeling [95, 96] or deconvolution [97]. Deconvolution can improve dynamic [26] and static FC estimates [97]. However, Seth et al. [27] have shown that sufficient sampling rate is more important for valid dynamic FC estimates. Unlike fMRI, electrophysiological measurements such as EEG and MEG have sufficient sampling rates and do not require deconvolution, so they could be used to study the relationship between static and dynamic FC [98]. Note that in EEG, volume conduction can inflate zero-lag connectivity, so careful consideration is needed to disentangle true zero-phase lag connectivity from volume conduction effects [99]. Furthermore, because instantaneous (zero-lag) signal transmission is not physiologically plausible, zero-phase lag effects in EEG most likely reflect indirect (non-causal) connections, whereas lagged effects are influenced by both indirect and direct (causal) connections. Therefore, the comparison of static and dynamic FC measures in the EEG can be used to disentangle causal and non-causal effects.

### 4.6. Conclusions

Our results show that the dynamic FC estimates represent information about connectivity that is broadly similar to the static FC. Moreover, we have shown that the similarity between dynamic and static FC is due, at least in part, to the convolution of neural time series with HRF. In contrast, we observed less similarity in the patterns of FC estimates between multivariate and bivariate methods. Multivariate FC methods were more sensitive to noise and CCA models based on multivariate methods were less generalizable. We also showed that the choice of FC methods affects the network topology, with noticable difference between multivariate and bivariate FC estimates, and only slight differences between dynamic and static FC estimates. While dynamic FC estimates can still provide information about the directionality of the connections, careful inspection of the results is required, as this information may change after averaging the FC matrices across participants.

Although dynamic FC models are useful as a model for directed FC or for modeling the evolution of neural time series over time [7], our results suggest that estimates of the functional connectome change very little when temporal information is taken into account. Dynamic FC estimates also show strong similarity to static FC in terms of brain-behavior associations.

## 5. Data and code availability

Raw data are available as part of the Human Connectome Project (https://www.humanconnectome.org/). The function to compute the variance component model is available in the repository: https://github.com/RaphaelLiegeois/FC-Behavior. For CCA and PLS, we used the GEMMR package: https://github.com/murraylab/gemmr. Strength- based node centrality measures were computed using the Brain Connectivity Toolbox (https://sites.google.com/site/bctnet/). The code for the normalized participation coefficient is available in the repository: https://github.com/omidvarnia/Dynamic_brain_connectivity_analysis. All other relevant code is available in the Open Science Framework repository: https://dx.doi.org/10.17605/OSF.IO/XFTDH.

## 6. Author contributions

**Andraž Matkovič:** Conceptualization, Methodology, Formal analysis, Investigation, Writing - Original Draft, Writing - Review & Editing, Visualization. **Alan Anticevic:** Conceptualization, Writing - Review & Editing. **John D. Murray:** Conceptualization, Writing - Review & Editing. **Grega Repovš:** Conceptualization, Software, Writing - Original Draft, Writing - Review & Editing, Supervision, Project administration, Funding acquisition.

## Acknowledgments

Data were provided by the Human Connectome Project, WU-Minn Consortium (Principal Investigators: David Van Essen and Kamil Ugurbil; 1U54MH091657) funded by the 16 NIH Institutes and Centers that support the NIH Blueprint for Neuroscience Research; and by the McDonnell Center for Systems Neuroscience at Washington University.

## 10. Supplement

**Figure S1:**
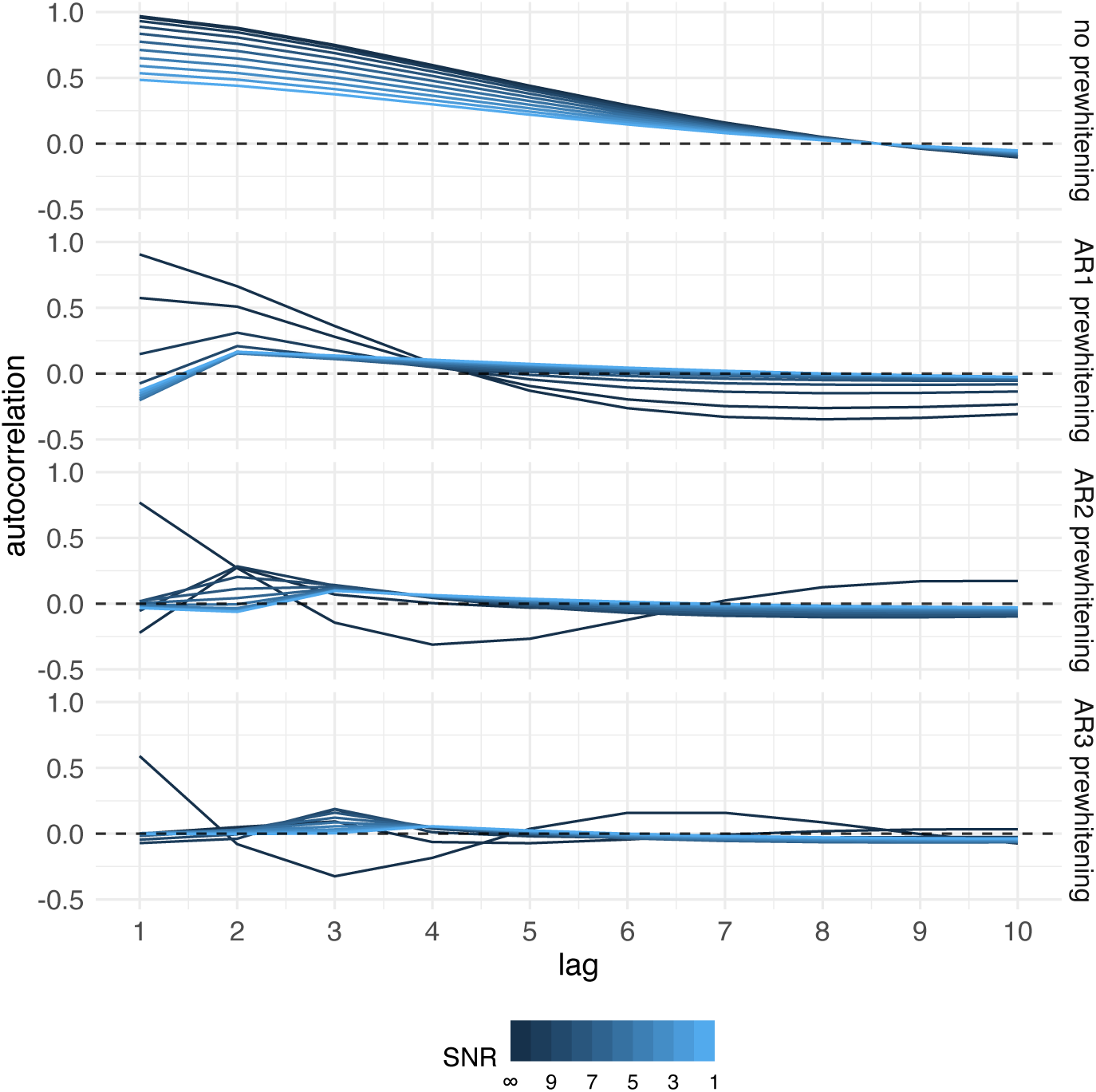
The autocorrelation function of simulated data as a function of prewhitening order and noise. The mean autocorrelation function was computed over all participants and regions. In general, noise and prewhitening reduced absolute autocorrelation. The shape of the autocorrelation function varied as a function of noise and prewhitening. In case without prewhitening, autocorrelation monotonically decreased and reached 0 at lag 8. After prewhitening, autocorrelation varied between positive and negative values, and this was most pronounced in cases without noise. The autocorrelation function was more similar to the experimental data in cases with low levels of noise.

**Figure S2:**
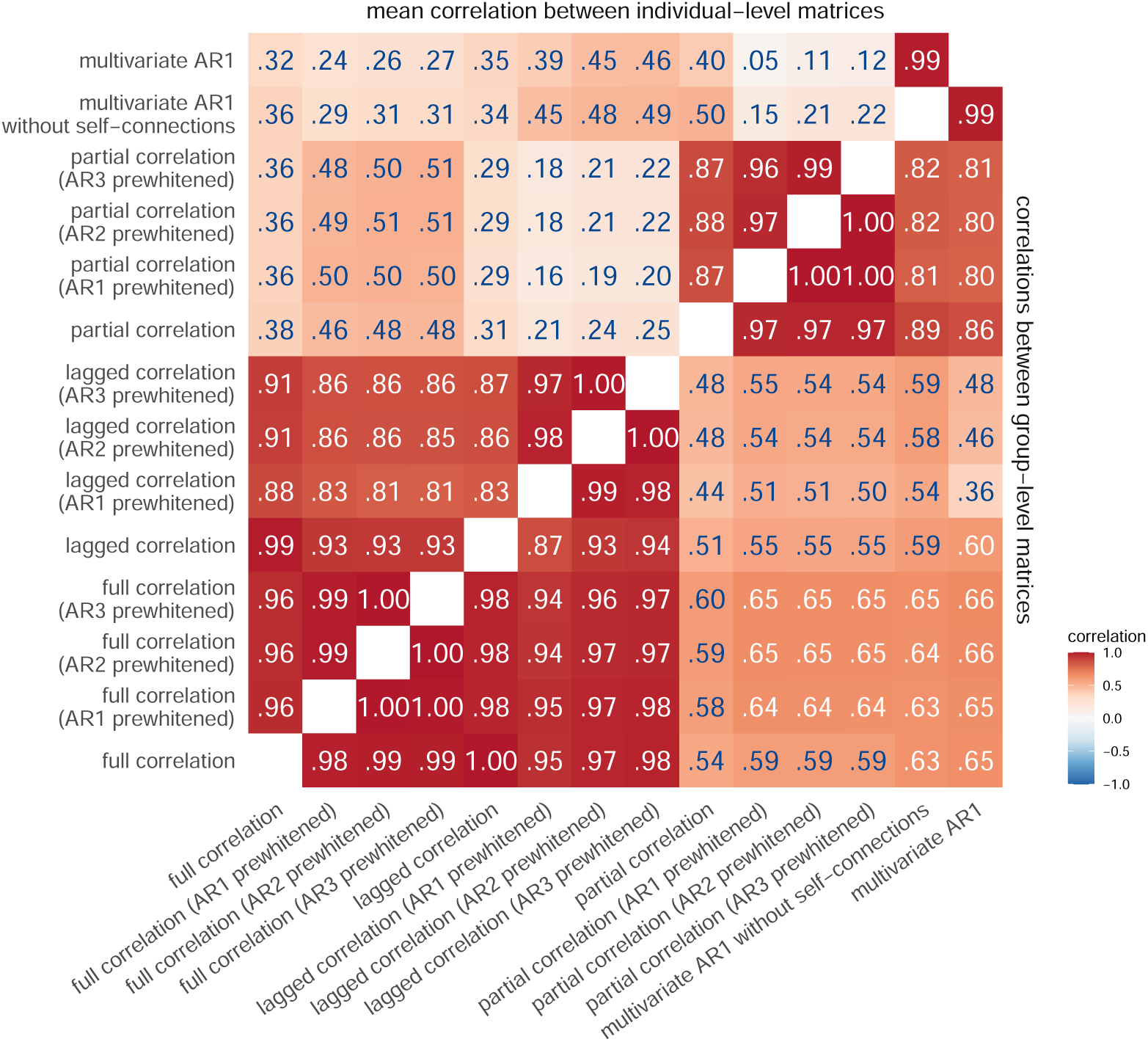
Correlations between connectivity methods. Same as in Figure 2A but includes all orders of prewhitening.

**Figure S3:**
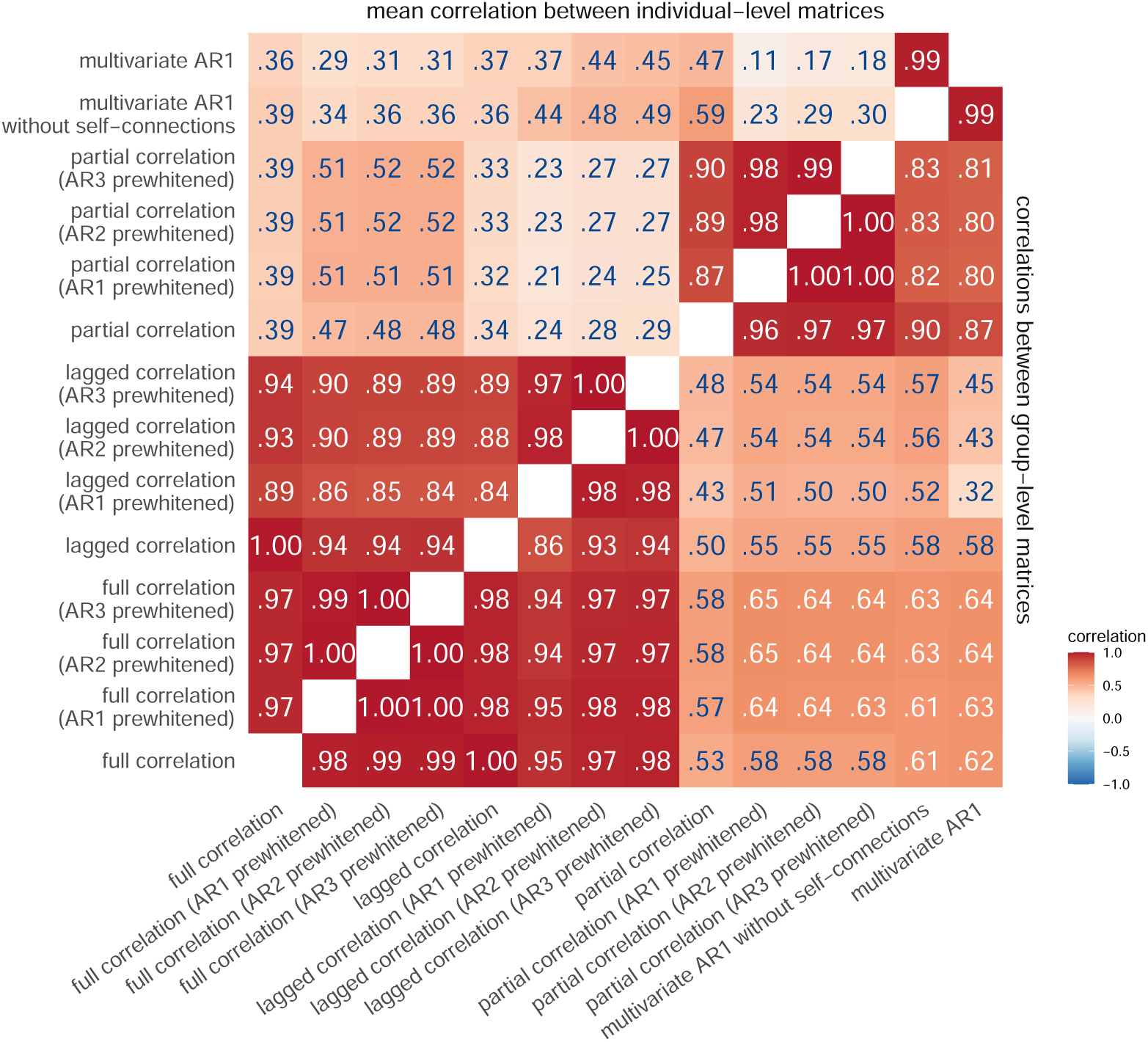
Correlations between connectivity methods on 200 participants with highest quality data.

**Figure S4:**
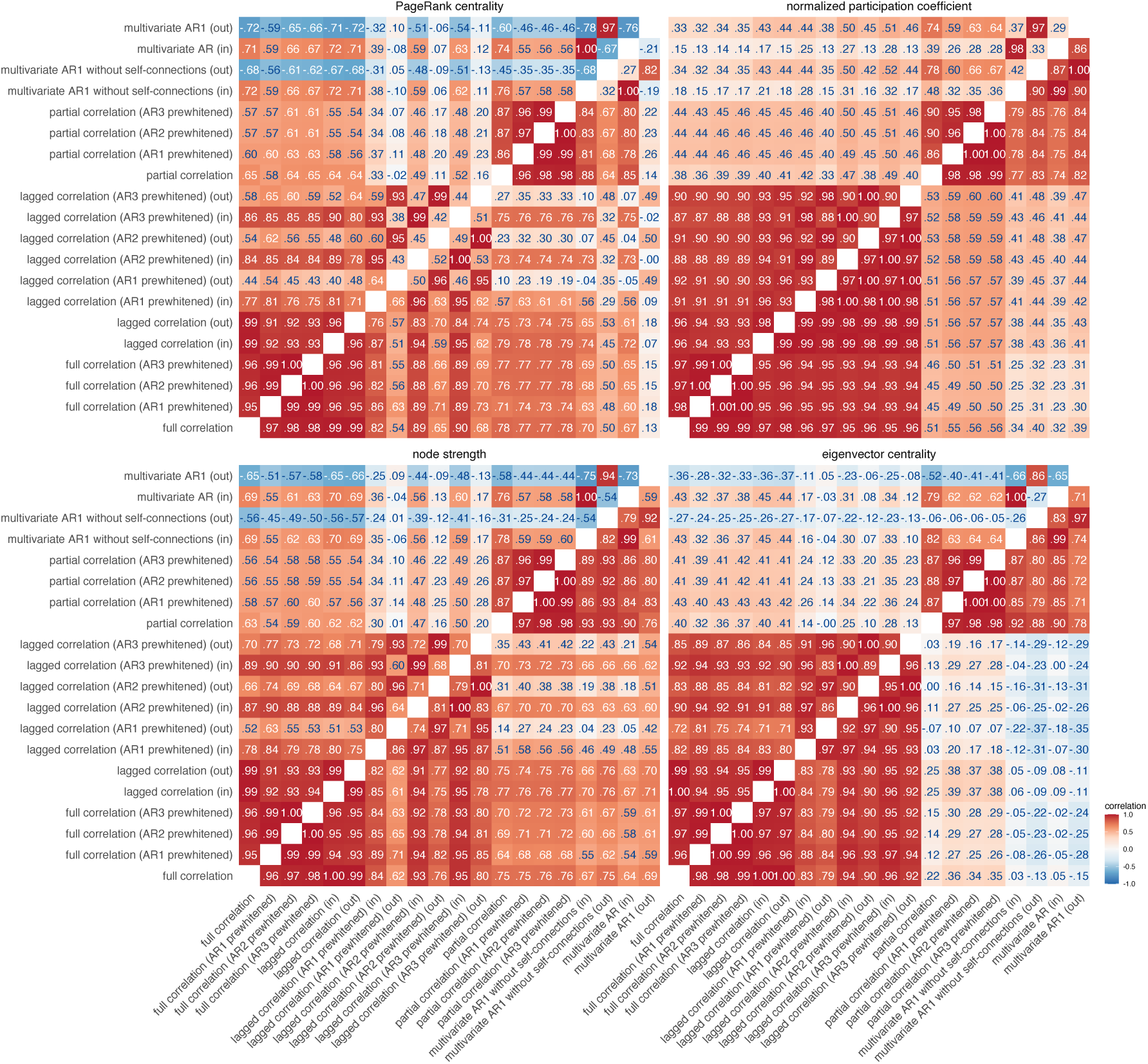
Similarities between node centrality measures based on positive connections. Similarities were estimated by (i) computing node measures on group-average connectivity matrices (group-level comparison; below diagonal), (ii) by computing node measures for each individual separately, correlating within participant and averaging these correlations across participants (individual-level comparsion; above diagonal). Same as in Figure 4, but includes prewhitened data.

**Figure S5:**
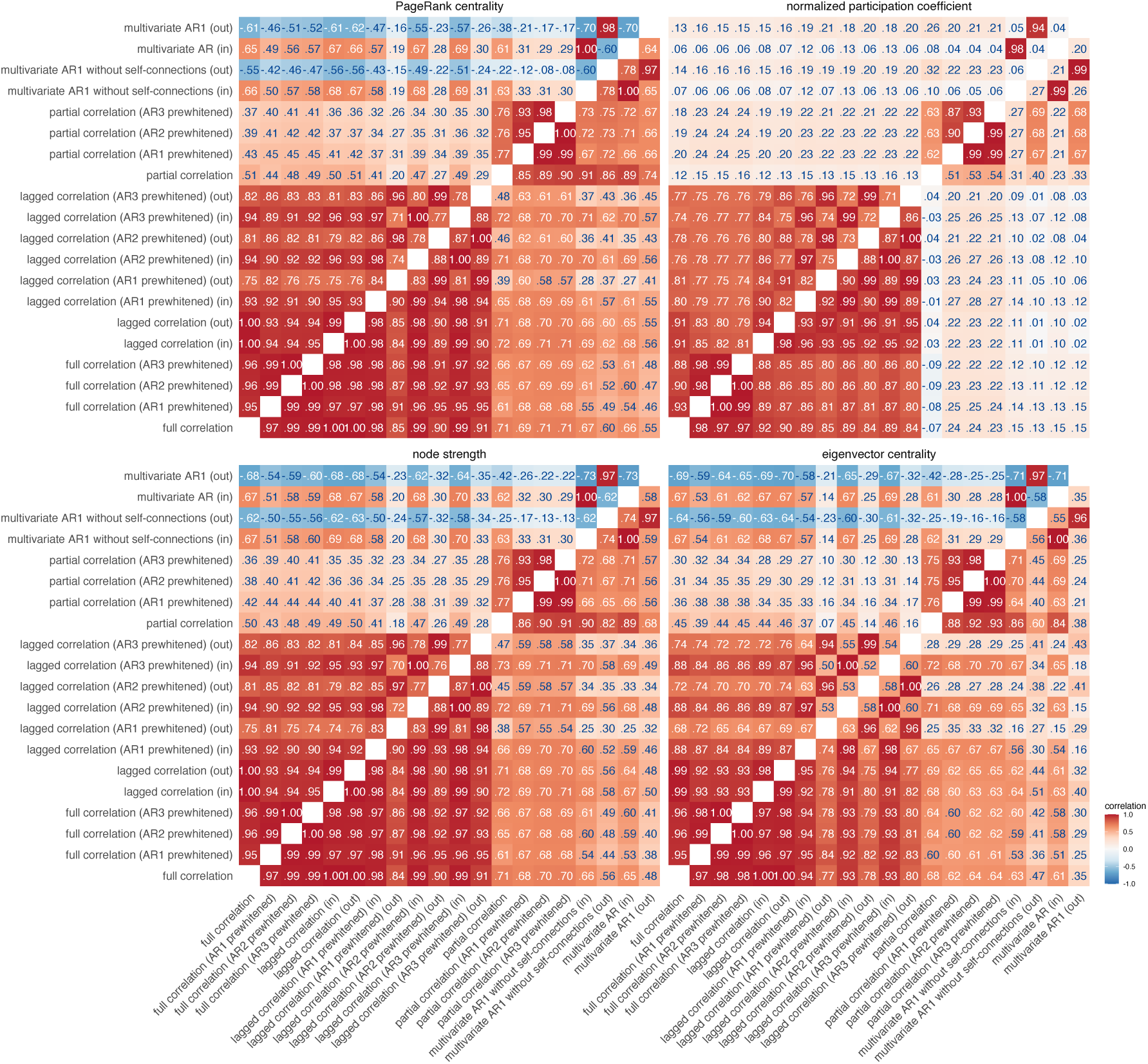
Similarities between node centrality measures based on positive connections. Similarities were estimated by (i) computing node measures on group-averaged connectivity matrices (group-level comparison; below diagonal), (ii) by computing node measures for each individual separately, correlating within participants and averaging these correlations across participants (individual-level comparison; above diagonal). Similar to Figure S4, but for negative connections.

**Figure S6:**
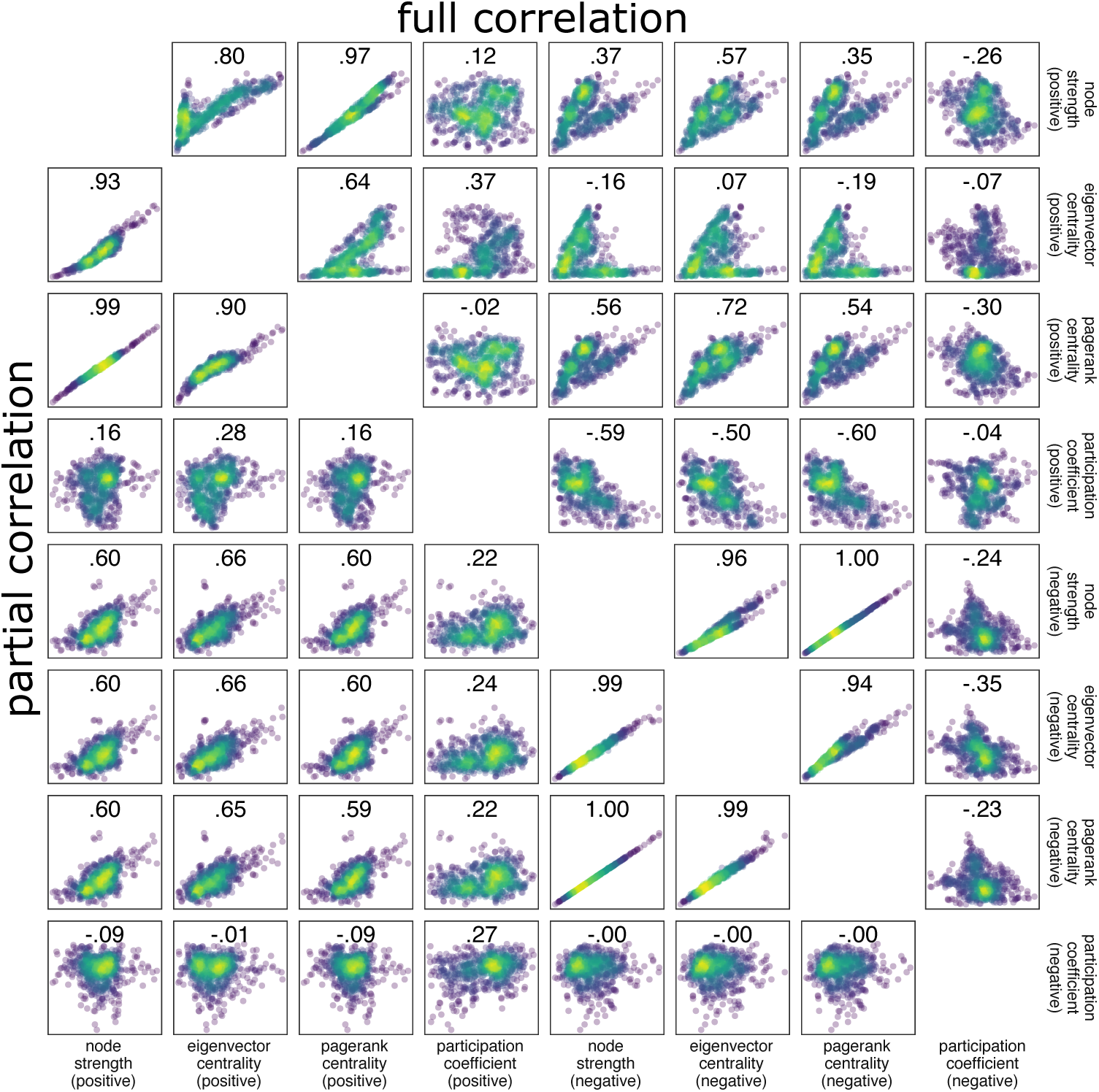
Correlations between centrality measures for static FC methods at the group level. Correlations were computed separately for positive and negative connections. We observed a positive correlation between the participation coefficient of positive connections and strength-based measures of negative connections. This suggests that nodes that participate in different modules tend to have fewer negative connections. Importantly, this finding highlights the functional importance of negative connections. However, for partial correlation networks, a positive correlation was found between strength-based measures and the participation coefficient. This suggests that indirect negative connections drive the negative relationship between participation coefficient and strength. In other words, nodes that participate in different modules tend to have more indirect negative functional connections, compared to nodes with low participation coefficient.

**Figure S7:**
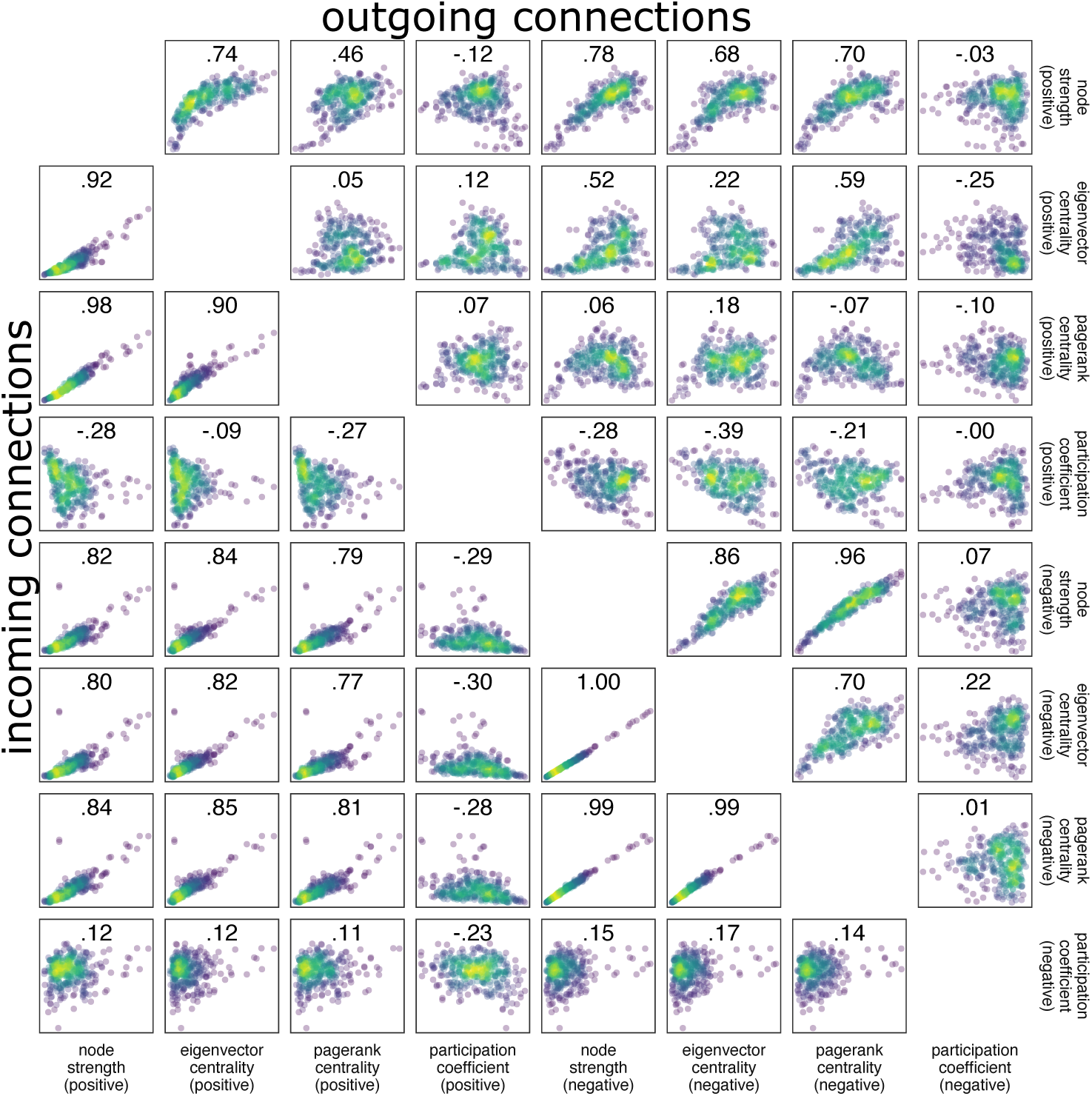
Correlations between centrality measures for the multivariate autoregressive model at the group level. Correlations were computed separately for positive and negative connections. The scatter plots above the diagonal refer to outgoing connections, while the scatter plots below the diagonal refer to incoming connections.

**Figure S8:**
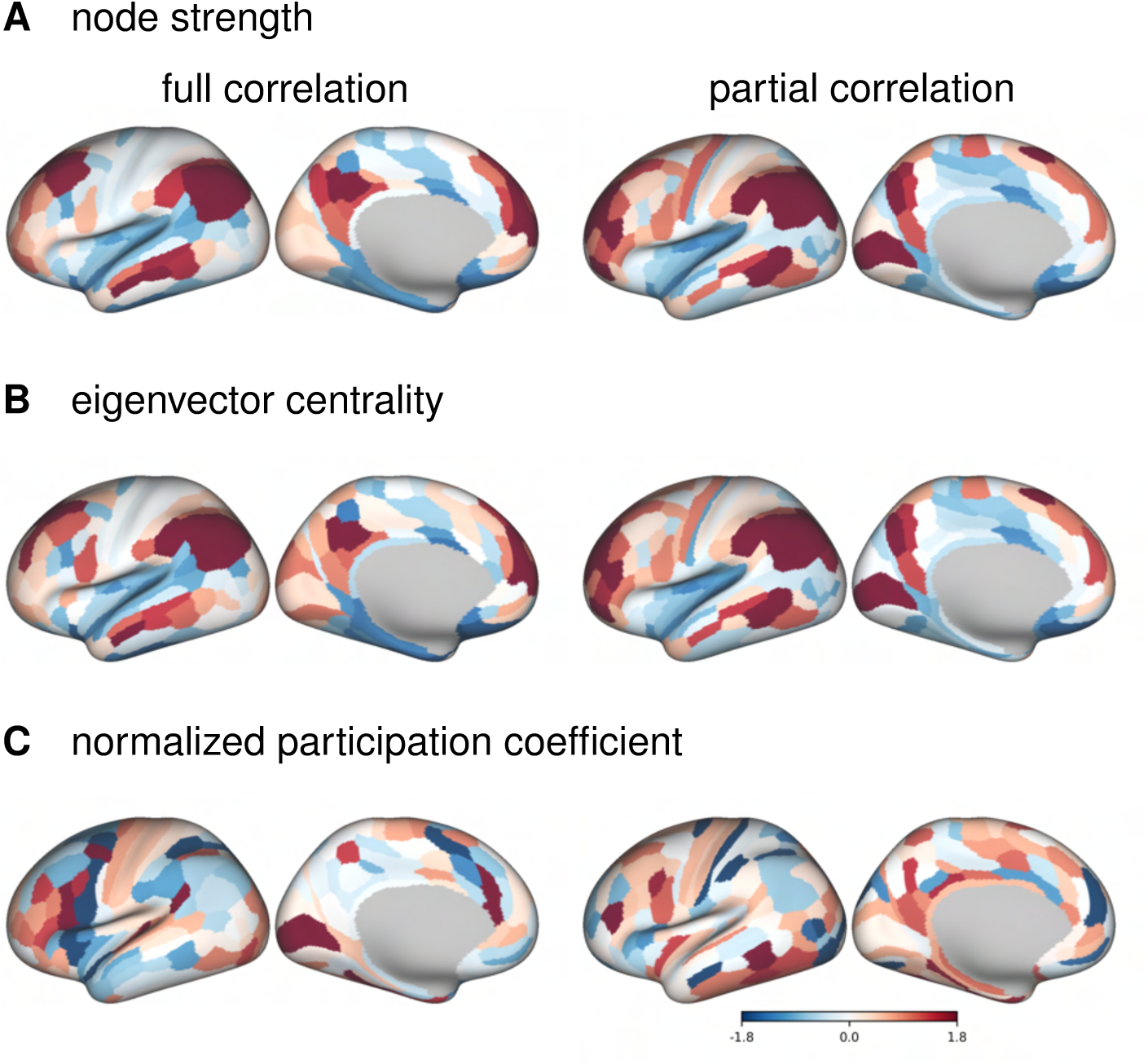
Cortical distribution of centrality measures for static FC methods and for negative connections. PageRank centrality is omitted, because its correlation with strength is equal to 1. The values have been transformed to *z*-values for visualization.

**Figure S9:**
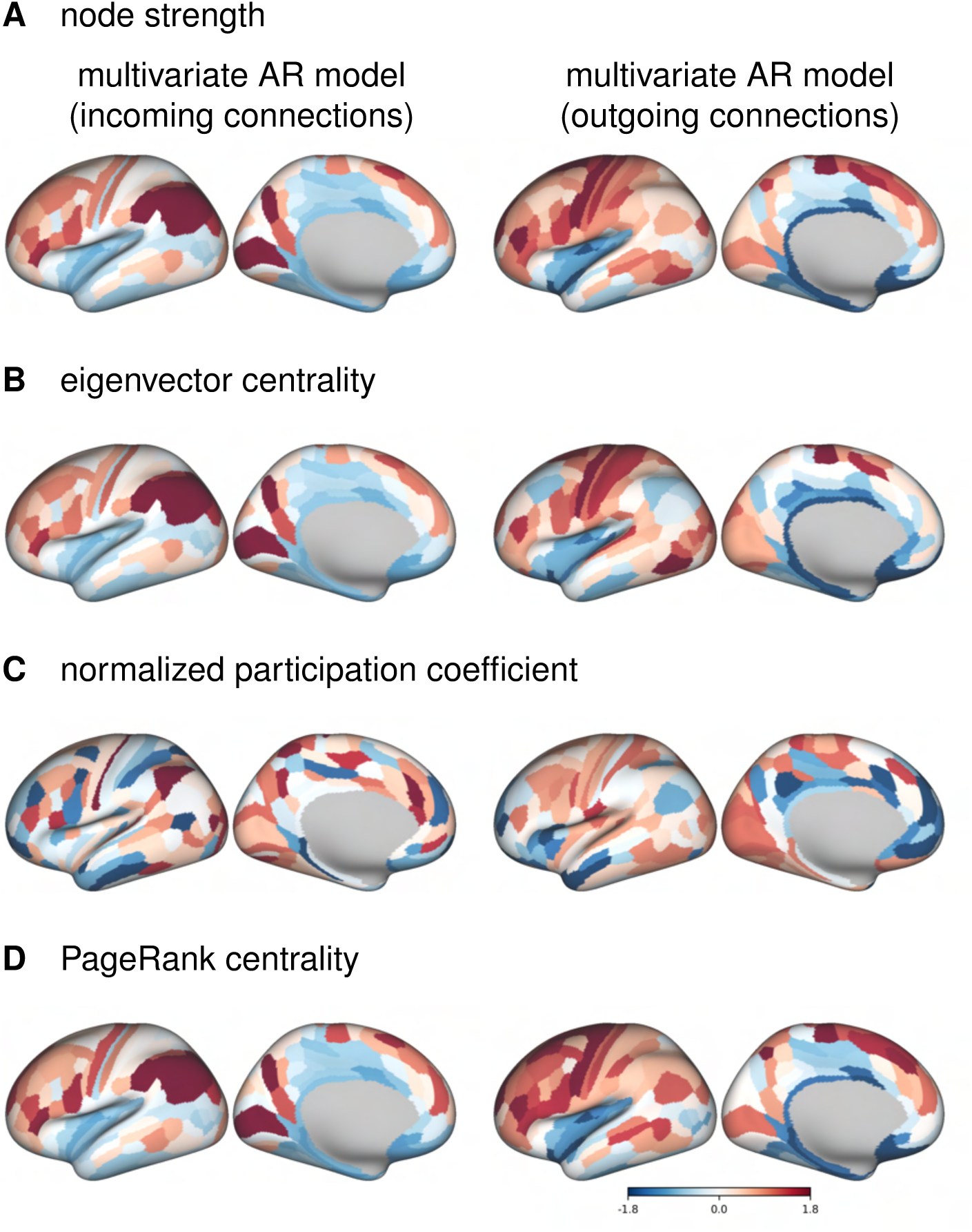
Cortical distribution of centrality measures for multivariate autoregressive model and for negative connections.

**Figure S10:**
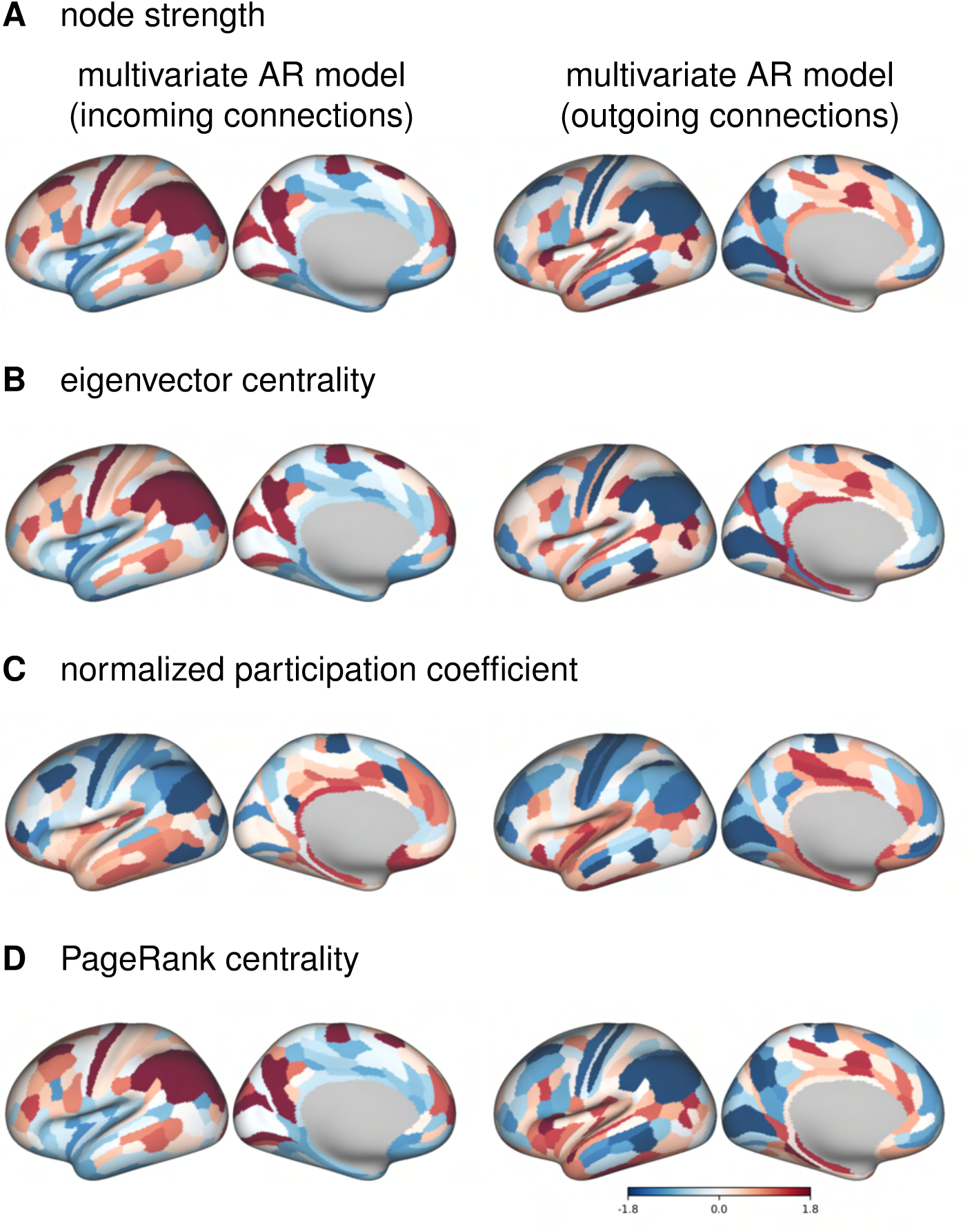
Cortical distribution of centrality measures for HCP subject 100307 for multivariate autoregressive model and for negative connections.

**Figure S11:**
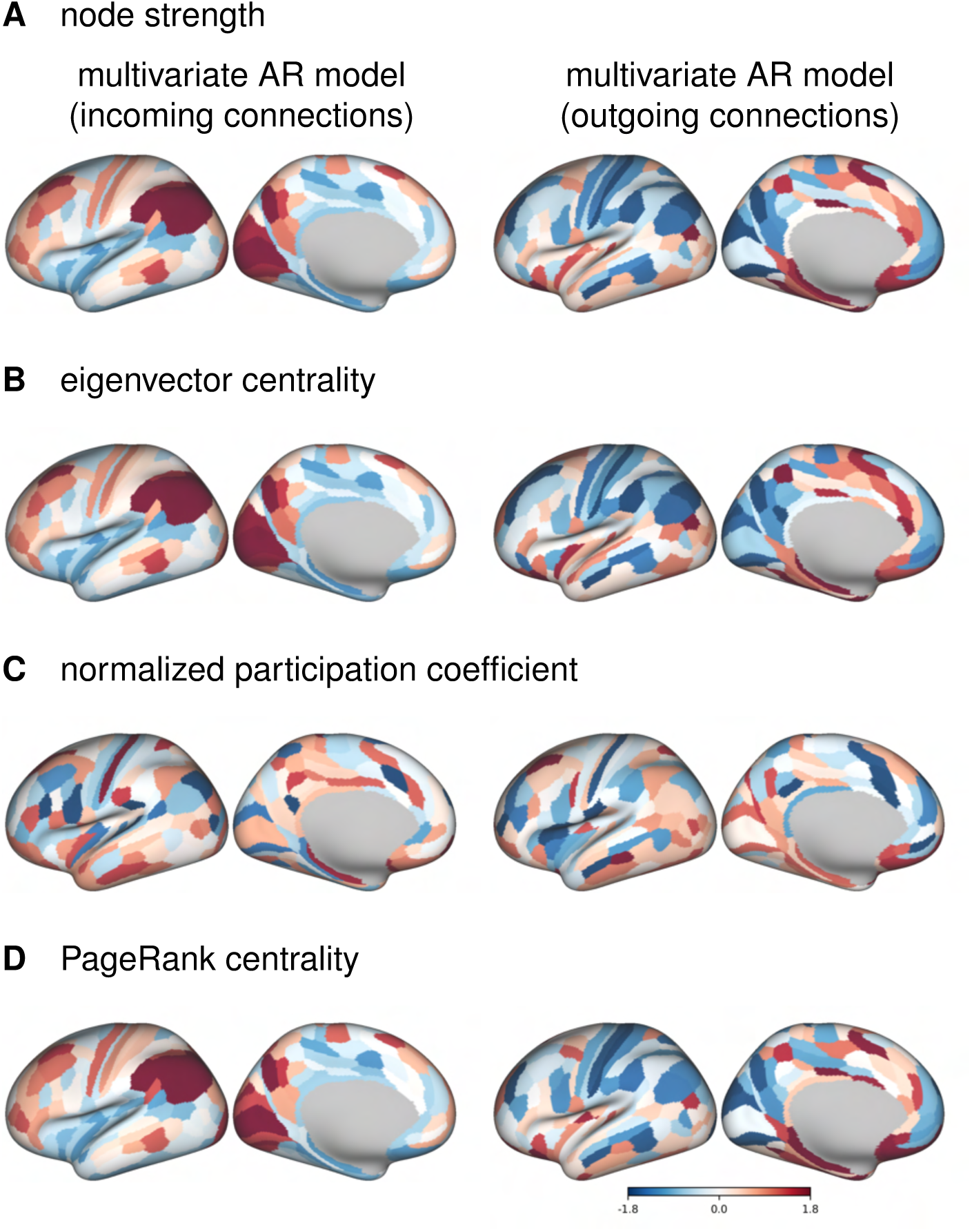
Cortical distribution of centrality measures for HCP subject 100307 for multivariate autoregressive model and for negative connections.

**Figure S12:**
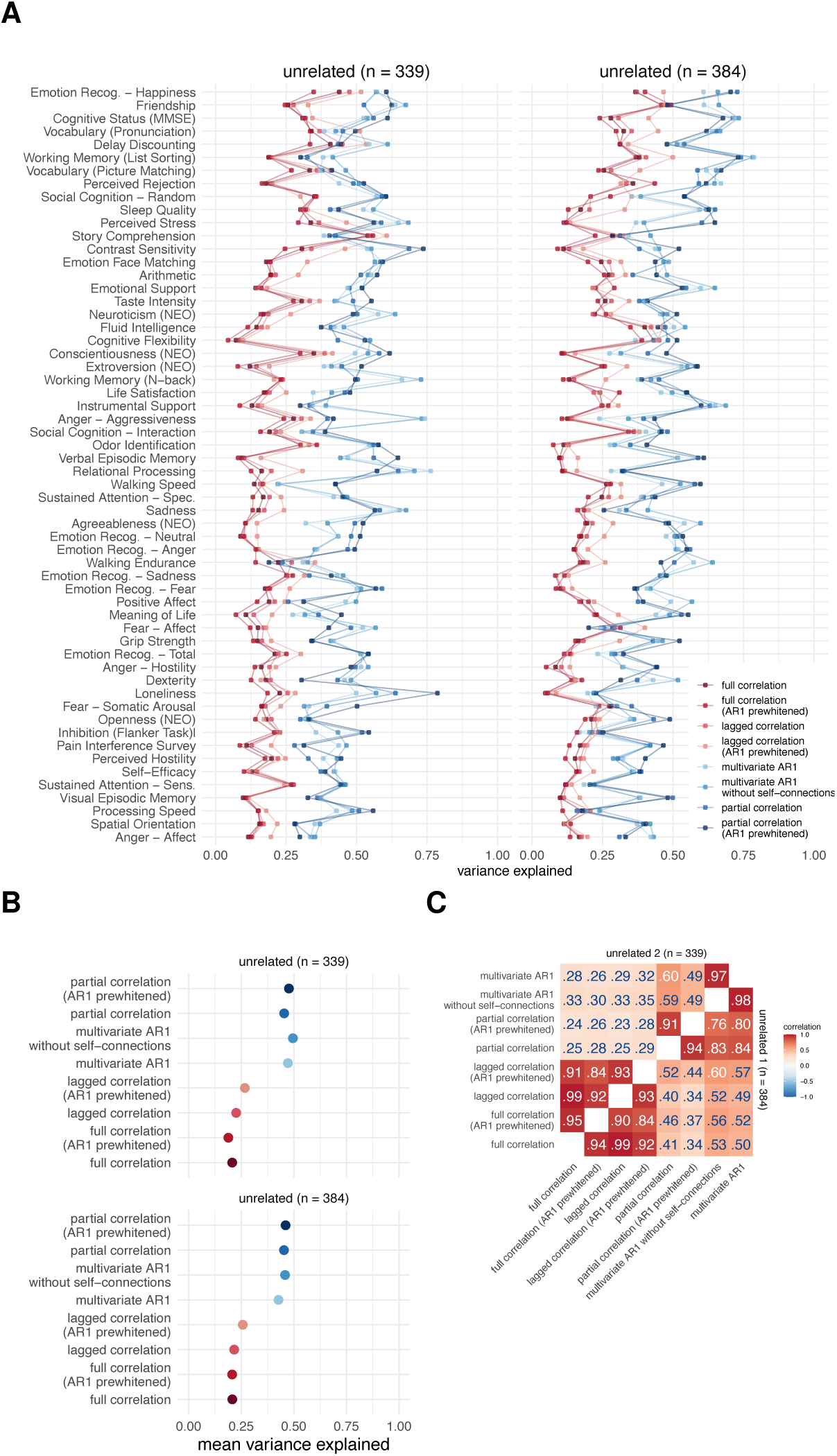
Results of variance component model for brain-behavior associations on subsamples of unrelated participants. (A) Variance explained for individual traits estimated with different connectivity methods, (B) mean variance explained, and (C) similarities of explained variance patterns between connectivity methods. The traits are ordered according to the mean variance explained across connectivity methods. The same as in Figure 7 but in subsamples of unrelated participants.

**Figure S13:**
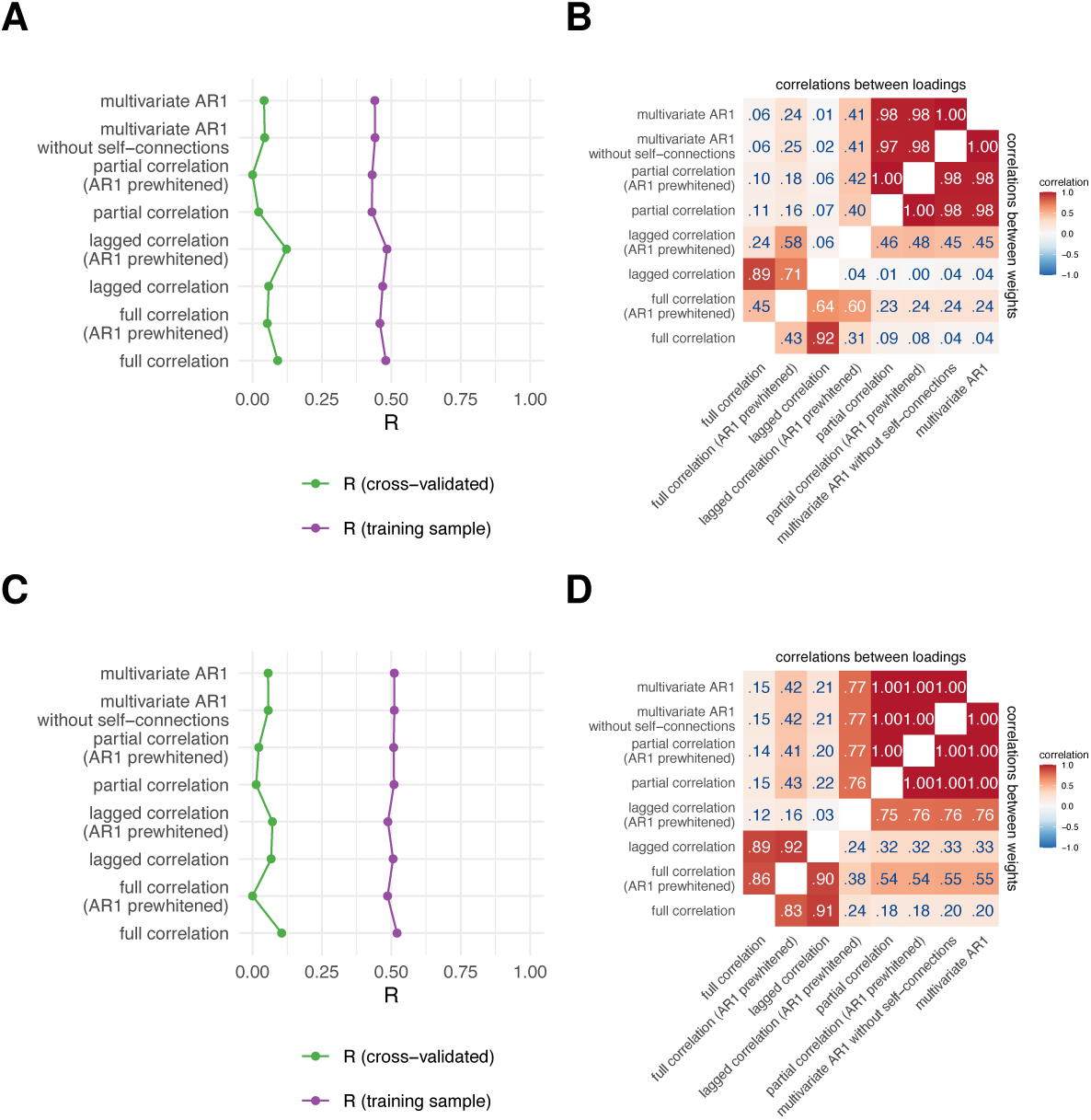
Results of canonical correlation analysis for brain-behavior associations on subsamples of unrelated participants. (A,C) First canonical correlation on test and training sets in the first (A, *n* = 384) and second subsample (C, *n* = 339). (B,D) Correlations between canonical loadings and weights across FC methods for the first canonical components on the first (B) and second (D) subsamples.

**Figure S14:**
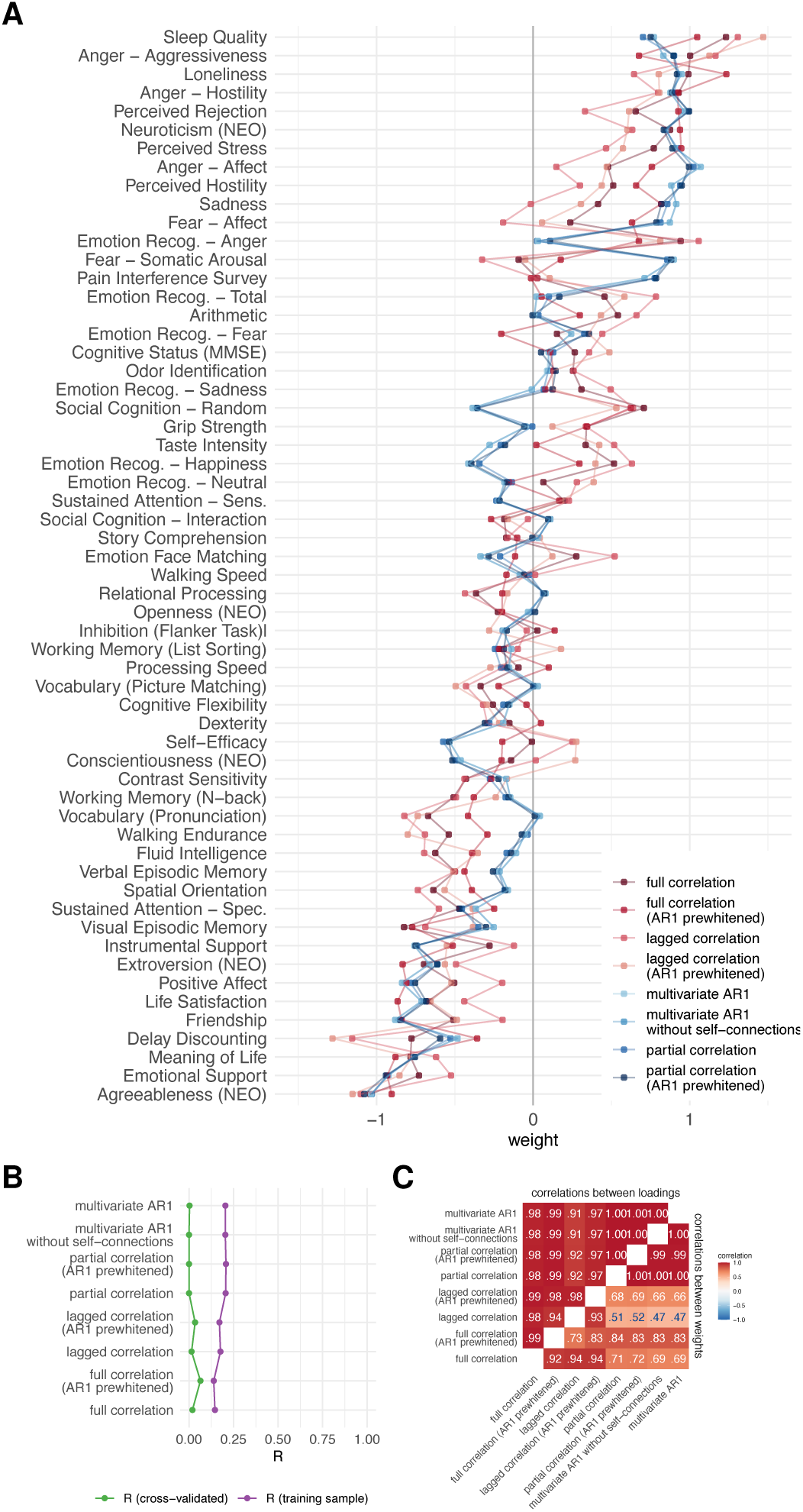
Results of principal least squares analysis for brain-behavior associations. **A.** PLS weights. **B.** First canonical correlation on test and training sets. **C.** Correlations between canonical loadings and weights across functional connectivity methods for first canonical components.

**Figure S15:**
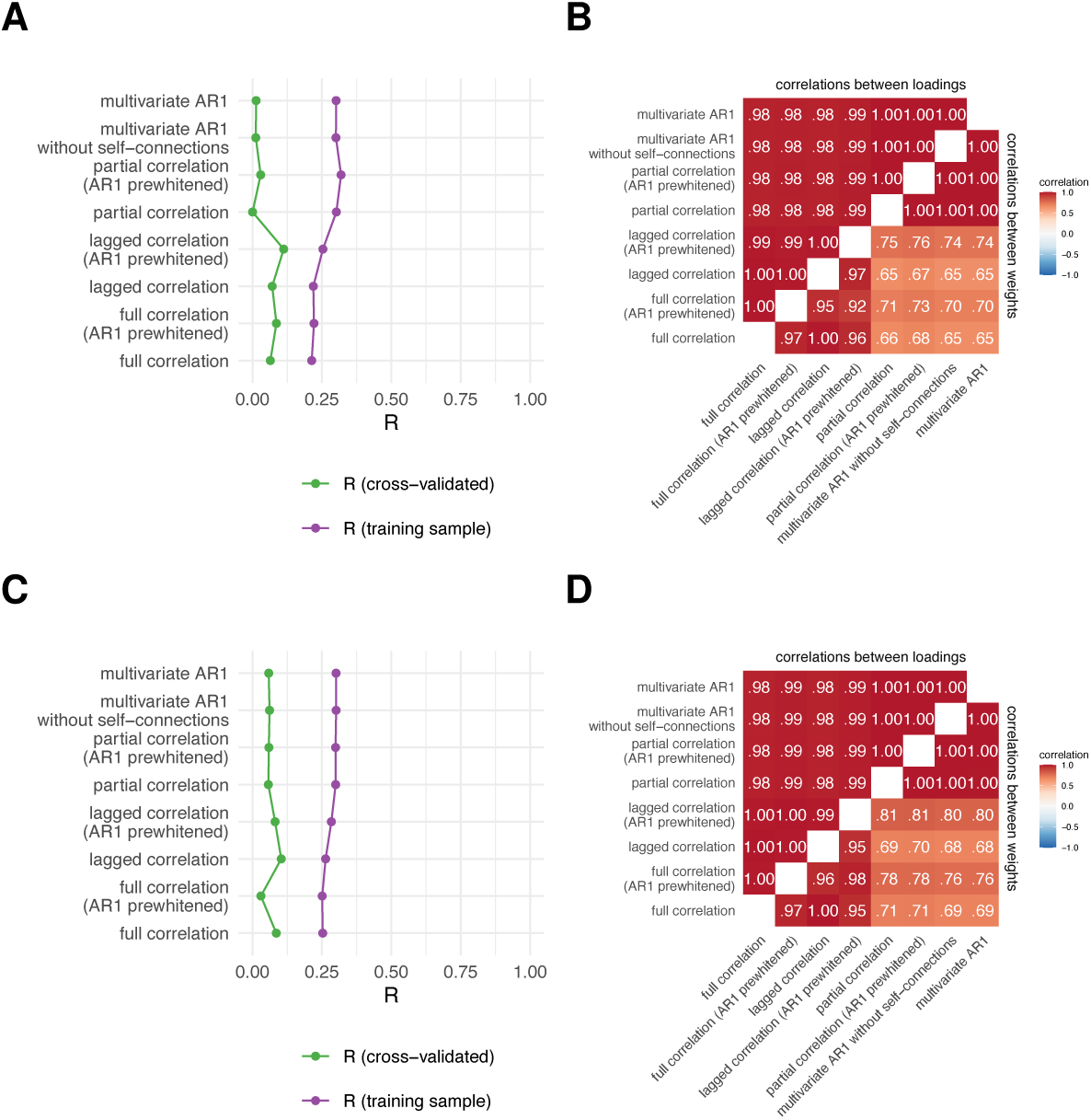
Results of principal least squares analysis for brain-behavior associations on subsamples of unrelated participants. (A,C) First canonical correlation on test and training sets in the first (A, *n* = 384) and second subsample (C, *n* = 339). (B,D) Correlations between canonical loadings and weights across FC methods for the first canonical components on the first (B) and second (D) subsamples.

**Figure S16:**
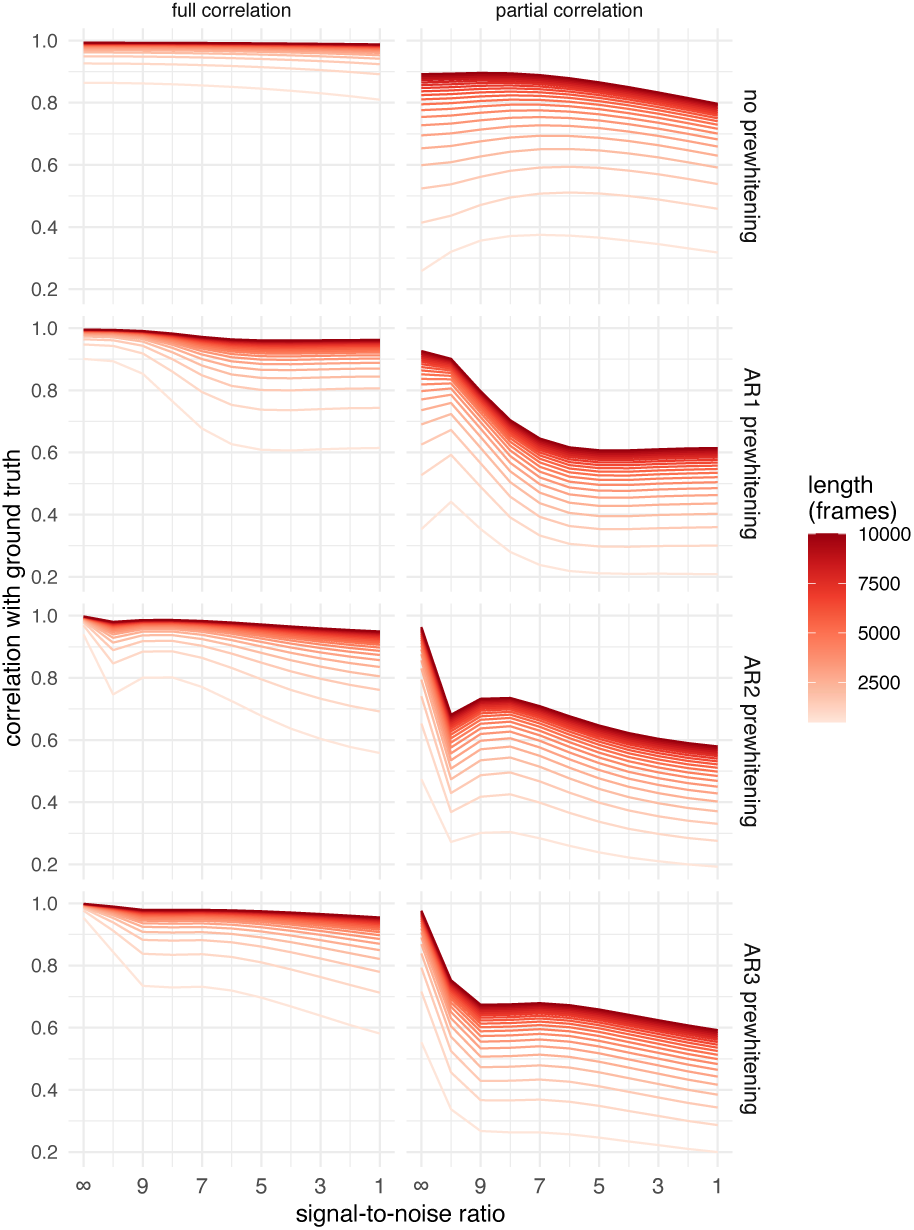
Correlation between ground truth and simulated data for all FC methods in association ith noise and signal length. Same as in Figure 11B but includes all orders of prewhitening.

**Figure S17:**
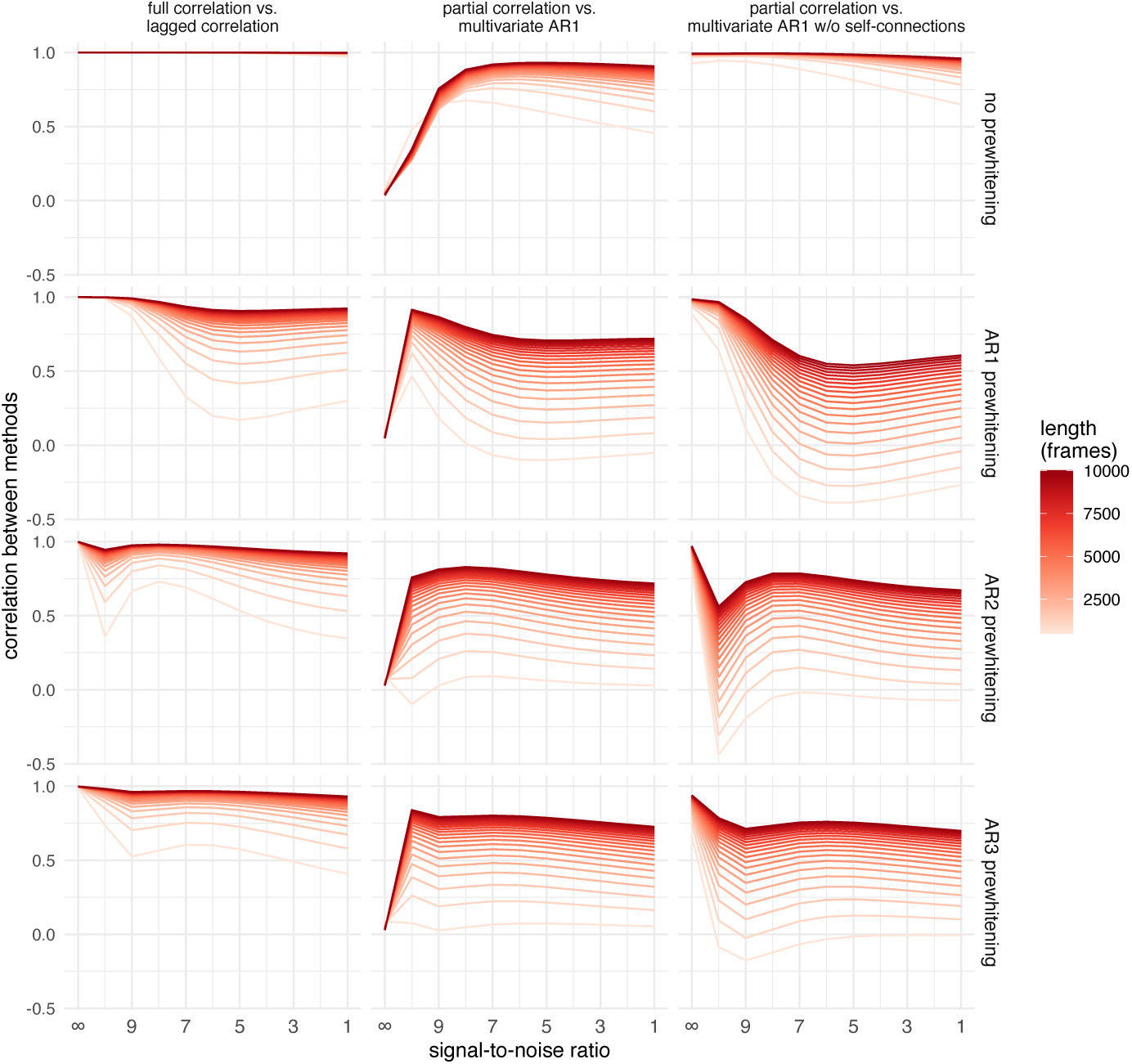
Correlation between selected pairs of FC methods as a function of noise and signal length on simulated data. Same as in Figure 11C but includes all prewhitening orders.

**Figure S18:**
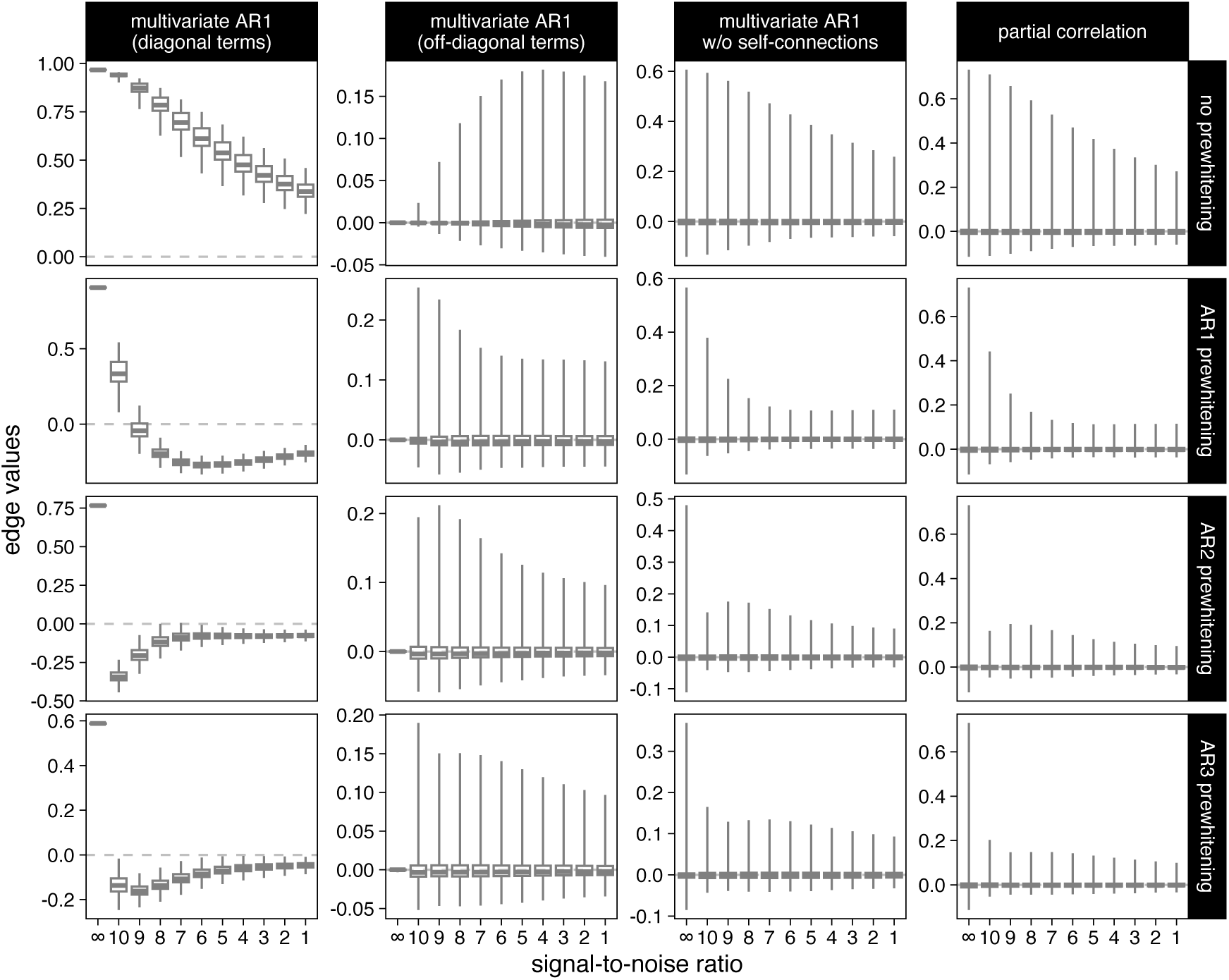
Distributions of edge values on simulated data for selected FC methods as a function of noise for the signals with the longest length (10000 frames). The distributions are based on the average FC matrix across simulated participants. The boxplot whiskers represent the minimum and maximum values.

**Table S1:**
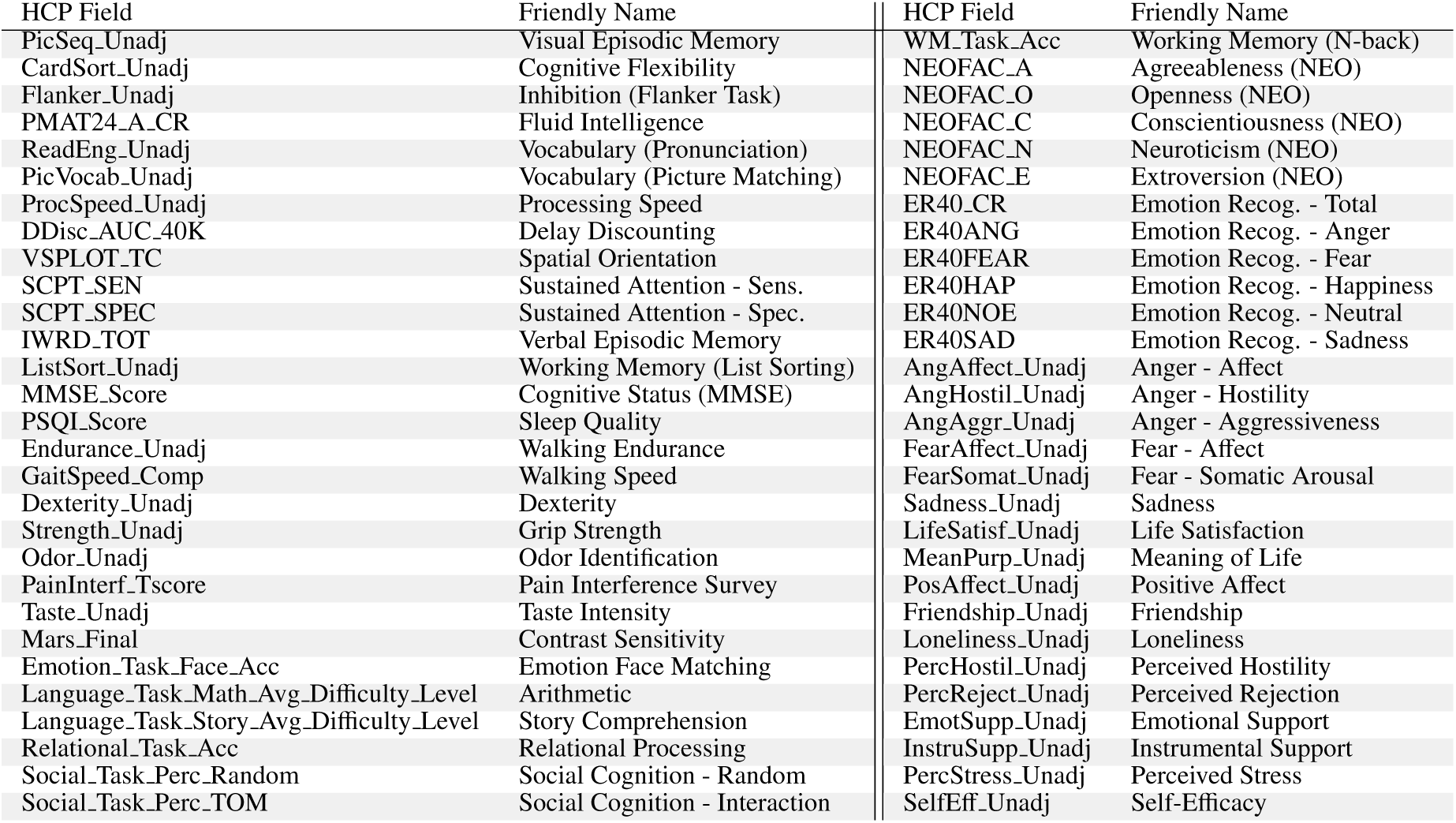
Behavioral measures.

## References

1. [1] A. Fornito, A. Zalesky, E. T. Bullmore, Fundamentals of Brain Network Analysis, Elsevier/Academic Press, Amsterdam ; Boston, 2016.

[2] A. T. Reid, D. B. Headley, R. D. Mill, R. Sanchez-Romero, L. Q. Uddin, D. Marinazzo, D. J. Lurie, P. A. Valdés-Sosa, S. J. Hanson, B. B. Biswal, V. Calhoun, R. A. Poldrack, M. W. Cole, Advancing functional connectivity research from association to causation, Nature Neuroscience 22 (2019) 1751–1760. doi:10.1038/s41593-019-0510-4.

[3] C. Wu, F. Ferreira, M. Fox, N. Harel, J. Hattangadi-Gluth, A. Horn, S. Jbabdi, J. Kahan, A. Oswal, S. A. Sheth, Y. Tie, V. Vakharia, L. Zrinzo, H. Akram, Clinical applications of magnetic resonance imaging based functional and structural connectivity, NeuroImage 244 (2021) 118649. doi:10.1016/j.neuroimage.2021.118649.

[4] S. H. Tompson, E. B. Falk, J. M. Vettel, D. S. Bassett, Network approaches to understand individual differences in brain connectivity: Opportunities for personality neuroscience, Personality Neuroscience 1 (2018) e5. doi:10.1017/pen.2018.4.

[5] M. W. Cole, T. Ito, D. S. Bassett, D. H. Schultz, Activity flow over resting-state networks shapes cognitive task activations, Nature Neuroscience 19 (2016) 1718–1726. doi:10.1038/nn.4406.

[6] T. A. W. Bolton, R. Liegeois, The arrow-of-time in neuroimaging time series identifies causal triggers of brain function, 2022. doi:10.1101/2022.05.11.491521.

[7] R. Liégeois, T. O. Laumann, A. Z. Snyder, J. Zhou, B. T. Yeo, Interpreting temporal fluctuations in resting-state functional connectivity MRI, NeuroImage 163 (2017) 437–455. doi:10/gcsbkz.

8. [8] C. Chatfield, H. Xing, The Analysis of Time Series: An Introduction with R, Chapman & Hall/CRC Texts in Statistical Science Series, seventh edition ed., CRC Press, Taylor & Francis Group, Boca Raton, 2019.

[9] M. G. Preti, T. A. Bolton, D. Van De Ville, The dynamic functional connectome: State-of-the-art and perspectives, NeuroImage 160 (2017) 41–54. doi:10/gcj5q7.

[10] D. J. Lurie, D. Kessler, D. S. Bassett, R. F. Betzel, M. Breakspear, S. Kheilholz, A. Kucyi, R. Liégeois, M. A. Lindquist, A. R. McIntosh, R. A. Poldrack, J. M. Shine, W. H. Thompson, N. Z. Bielczyk, L. Douw, D. Kraft, R. L. Miller, M. Muthuraman, L. Pasquini, A. Razi, D. Vidaurre, H. Xie, V. D. Calhoun, Questions and controversies in the study of time-varying functional connectivity in resting fMRI, Network Neuroscience 4 (2020) 30–69. doi:10/ghr3sd.

[11] R. Liégeois, B. T. Yeo, D. Van De Ville, Interpreting null models of resting-state functional MRI dynamics: Not throwing the model out with the hypothesis, NeuroImage 243 (2021) 118518. doi:10.1016/j.neuroimage.2021.118518.

[12] R. Hindriks, M. Adhikari, Y. Murayama, M. Ganzetti, D. Mantini, N. Logothetis, G. Deco, Can sliding-window correlations reveal dynamic functional connectivity in resting-state fMRI?, NeuroImage 127 (2016) 242–256. doi:10/f786fz.

[13] T. O. Laumann, A. Z. Snyder, A. Mitra, E. M. Gordon, C. Gratton, B. Adeyemo, A. W. Gilmore, S. M. Nelson, J. J. Berg, D. J. Greene, J. E. McCarthy, E. Tagliazucchi, H. Laufs, B. L. Schlaggar, N. U. F. Dosenbach, S. E. Petersen, On the stability of bold fmri correlations, Cerebral Cortex (2016) cercor;bhw265v1. doi:10/f9p8v2.

[14] J. Daniel Arzate-Mena, E. Abela, P. V. Olguín-Rodríguez, W. Ríos-Herrera, S. Alcauter, K. Schindler, R. Wiest, M. F. Müller, C. Rummel, Stationary EEG pattern relates to large-scale resting state networks – An EEG- fMRI study connecting brain networks across time-scales, NeuroImage 246 (2022) 118763. doi:10.1016/j.neuroimage.2021.118763.

15. P. V. Olguín-Rodríguez, J. D. Arzate-Mena, M. Corsi-Cabrera, H. Gast, A. Marín-García, J. Mathis, J. Ramos Loyo, I. Y. Del Rio-Portilla, C. Rummel, K. Schindler, M. Müller, Characteristic fluctuations around stable attractor dynamics extracted from highly nonstationary electroencephalographic recordings, Brain Connectivity 8 (2018) 457–474. doi:10.1089/brain.2018.0609.

[16] M. F. Müller, C. Rummel, M. Goodfellow, K. Schindler, Standing waves as an explanation for generic stationary correlation patterns in noninvasive EEG of focal onset seizures, Brain Connectivity 4 (2014) 131–144. doi:10.1089/brain.2013.0192.

[17] D. A. Handwerker, V. Roopchansingh, J. Gonzalez-Castillo, P. A. Bandettini, Periodic changes in fMRI connectivity, NeuroImage 63 (2012) 1712–1719. doi:10.1016/j.neuroimage.2012.06.078.

[18] C. Chang, G. H. Glover, Time–frequency dynamics of resting-state brain connectivity measured with fMRI, NeuroImage 50 (2010) 81–98. doi:10.1016/j.neuroimage.2009.12.011.

[19] A. Zalesky, A. Fornito, L. Cocchi, L. L. Gollo, M. Breakspear, Time-resolved resting-state brain networks, Proceedings of the National Academy of Sciences 111 (2014) 10341–10346. doi:10.1073/pnas.1400181111.

[20] L. Novelli, A. Razi, A mathematical perspective on edge-centric brain functional connectivity, Nature Communications 13 (2022) 2693. doi:10.1038/s41467-022-29775-7.

[21] Z. Ladwig, B. A. Seitzman, A. Dworetsky, Y. Yu, B. Adeyemo, D. M. Smith, S. E. Petersen, C. Gratton, BOLD cofluctuation ‘events’ are predicted from static functional connectivity, NeuroImage 260 (2022) 119476. doi:10.1016/j.neuroimage.2022.119476.

[22] T. Matsui, T. Q. Pham, K. Jimura, J. Chikazoe, On co-activation pattern analysis and non-stationarity of resting brain activity, NeuroImage 249 (2022) 118904. doi:10.1016/j.neuroimage.2022.118904.

23. S. M. Smith, K. L. Miller, G. Salimi-Khorshidi, M. Webster, C. F. Beckmann, T. E. Nichols, J. D. Ramsey, M. W. Woolrich, Network modelling methods for FMRI, NeuroImage 54 (2011) 875–891. doi:10.1016/j.neuroimage.2010.08.063.

[24] K. Friston, Causal modelling and brain connectivity in functional magnetic resonance imaging, PLoS Biology 7 (2009) e1000033. doi:10.1371/journal.pbio.1000033.

[25] K. Friston, Dynamic causal modeling and Granger causality Comments on: The identification of interacting networks in the brain using fMRI: Model selection, causality and deconvolution, NeuroImage 58 (2011) 303–305. doi:10.1016/j.neuroimage.2009.09.031.

[26] O. David, I. Guillemain, S. Saillet, S. Reyt, C. Deransart, C. Segebarth, A. Depaulis, Identifying neural drivers with functional mri: An electrophysiological validation, PLoS Biology 6 (2008) e315. doi:10.1371/journal.pbio.0060315.

[27] A. K. Seth, P. Chorley, L. C. Barnett, Granger causality analysis of fMRI BOLD signals is invariant to hemodynamic convolution but not downsampling, NeuroImage 65 (2013) 540–555. doi:10.1016/j.neuroimage.2012.09.049.

[28] V. Pallarés, A. Insabato, A. Sanjuán, S. Kühn, D. Mantini, G. Deco, M. Gilson, Extracting orthogonal subject- and condition-specific signatures from fMRI data using whole-brain effective connectivity, NeuroImage 178 (2018) 238–254. doi:10.1016/j.neuroimage.2018.04.070.

[29] M. Gilson, G. Deco, K. J. Friston, P. Hagmann, D. Mantini, V. Betti, G. L. Romani, M. Corbetta, Effective connectivity inferred from fMRI transition dynamics during movie viewing points to a balanced reconfiguration of cortical interactions, NeuroImage 180 (2018) 534–546. doi:10.1016/j.neuroimage.2017.09.061.

[30] M. Gilson, R. Moreno-Bote, A. Ponce-Alvarez, P. Ritter, G. Deco, Estimation of directed effective connectivity from fmri functional connectivity hints at asymmetries of cortical connectome, PLOS Computational Biology 12 (2016) e1004762. doi:10.1371/journal.pcbi.1004762.

[31] C. Jin, H. Jia, P. Lanka, D. Rangaprakash, L. Li, T. Liu, X. Hu, G. Deshpande, Dynamic brain connectivity is a better predictor of PTSD than static connectivity: Dynamic Brain Connectivity, Human Brain Mapping 38 (2017) 4479–4496. doi:10.1002/hbm.23676.

[32] R. Liégeois, J. Li, R. Kong, C. Orban, D. Van De Ville, T. Ge, M. R. Sabuncu, B. T. T. Yeo, Resting brain dynamics at different timescales capture distinct aspects of human behavior, Nature Communications 10 (2019) 2317. doi:10/gf3k2q.

[33] M. R. Arbabshirani, A. Preda, J. G. Vaidya, S. G. Potkin, G. Pearlson, J. Voyvodic, D. Mathalon, T. van Erp, A. Michael, K. A. Kiehl, J. A. Turner, V. D. Calhoun, Autoconnectivity: A new perspective on human brain function, Journal of Neuroscience Methods 323 (2019) 68–76. doi:10/gf2tvj.

[34] R. Liégeois, A. Santos, V. Matta, D. Van De Ville, A. H. Sayed, Revisiting correlation-based functional connectivity and its relationship with structural connectivity, Network Neuroscience 4 (2020) 1235–1251. doi:10/gm79q6.

[35] H. Honari, A. S. Choe, J. J. Pekar, M. A. Lindquist, Investigating the impact of autocorrelation on time-varying connectivity, NeuroImage 197 (2019) 37–48. doi:10/gm7955.

[36] L. Novelli, J. T. Lizier, Inferring network properties from time series using transfer entropy and mutual information: Validation of multivariate versus bivariate approaches, Network Neuroscience (2021) 1–32. doi:10.1162/netn_a_00178.

[37] W. Cheng, X. Ji, J. Zhang, J. Feng, Individual classification of ADHD patients by integrating multiscale neuroimaging markers and advanced pattern recognition techniques, Frontiers in Systems Neuroscience 6 (2012). doi:10.3389/fnsys.2012.00058.

[38] H. Cai, J. Zhu, Y. Yu, Robust prediction of individual personality from brain functional connectome, Social Cognitive and Affective Neuroscience 15 (2020) 359–369. doi:10.1093/scan/nsaa044.

[39] A. Abraham, M. P. Milham, A. Di Martino, R. C. Craddock, D. Samaras, B. Thirion, G. Varoquaux, Deriving reproducible biomarkers from multi-site resting-state data: An Autism-based example, NeuroImage 147 (2017) 736–745. doi:10.1016/j.neuroimage.2016.10.045.

[40] K. Dadi, M. Rahim, A. Abraham, D. Chyzhyk, M. Milham, B. Thirion, G. Varoquaux, Benchmarking functional connectome-based predictive models for resting-state fMRI, NeuroImage 192 (2019) 115–134. doi:10.1016/j.neuroimage.2019.02.062.

[41] A. Zalesky, A. Fornito, E. Bullmore, On the use of correlation as a measure of network connectivity, NeuroImage 60 (2012) 2096–2106. doi:10.1016/j.neuroimage.2012.02.001.

[42] X. Liang, J. Wang, C. Yan, N. Shu, K. Xu, G. Gong, Y. He, Effects of different correlation metrics and preprocessing factors on small-world brain functional networks: A resting-state functional MRI study, PLoS ONE 7 (2012) e32766. doi:10.1371/journal.pone.0032766.

[43] D. C. Van Essen, S. M. Smith, D. M. Barch, T. E. Behrens, E. Yacoub, K. Ugurbil, The WU-Minn Human Connectome Project: An overview, NeuroImage 80 (2013) 62–79. doi:10/f46ktq.

[44] M. F. Glasser, S. N. Sotiropoulos, J. A. Wilson, T. S. Coalson, B. Fischl, J. L. Andersson, J. Xu, S. Jbabdi, M. Webster, J. R. Polimeni, D. C. Van Essen, M. Jenkinson, The minimal preprocessing pipelines for the Human Connectome Project, NeuroImage 80 (2013) 105–124. doi:10/f46nj4.

[45] G. Salimi-Khorshidi, G. Douaud, C. F. Beckmann, M. F. Glasser, L. Griffanti, S. M. Smith, Automatic denoising of functional MRI data: Combining independent component analysis and hierarchical fusion of classifiers, NeuroImage 90 (2014) 449–468. doi:10/ggwbcj.

[46] E. C. Robinson, S. Jbabdi, M. F. Glasser, J. Andersson, G. C. Burgess, M. P. Harms, S. M. Smith, D. C. Van Essen, M. Jenkinson, MSM: A new flexible framework for Multimodal Surface Matching, NeuroImage 100 (2014) 414–426. doi:10.1016/j.neuroimage.2014.05.069.

[47] J. L. Ji, J. Demšar, C. Fonteneau, Z. Tamayo, L. Pan, A. Kraljič, A. Matkovič, N. Purg, M. Helmer, S. Warrington, A. Winkler, V. Zerbi, T. S. Coalson, M. F. Glasser, M. P. Harms, S. N. Sotiropoulos, J. D. Murray, A. Anticevic, G. Repovš, QuNex—An integrative platform for reproducible neuroimaging analytics, Frontiers in Neuroinformatics 17 (2023) 1104508. doi:10.3389/fninf.2023.1104508.

[48] M. R. Arbabshirani, E. Damaraju, R. Phlypo, S. Plis, E. Allen, S. Ma, D. Mathalon, A. Preda, J. G. Vaidya, T. Adali, V. D. Calhoun, Impact of autocorrelation on functional connectivity, NeuroImage 102 (2014) 294–308. doi:10/f6rcdt.

[49] C. E. Davey, D. B. Grayden, G. F. Egan, L. A. Johnston, Filtering induces correlation in fMRI resting state data, NeuroImage 64 (2013) 728–740. doi:10/f4jgxv.

[50] M. F. Glasser, T. S. Coalson, E. C. Robinson, C. D. Hacker, J. Harwell, E. Yacoub, K. Ugurbil, J. Andersson, C. F. Beckmann, M. Jenkinson, S. M. Smith, D. C. Van Essen, A multi-modal parcellation of human cerebral cortex, Nature 536 (2016) 171–178. doi:10/f8z3gb.

[51] U. Pervaiz, D. Vidaurre, M. W. Woolrich, S. M. Smith, Optimising network modelling methods for fMRI, NeuroImage 211 (2020) 116604. doi:10/ggx68f.

[52] J. Friedman, T. Hastie, R. Tibshirani, Regularization paths for generalized linear models via coordinate descent, Journal of Statistical Software 33 (2010). doi:10.18637/jss.v033.i01.

[53] C.-M. Ting, A.-K. Seghouane, M. U. Khalid, S.-H. Salleh, Is first-order vector autoregressive model optimal for fMRI data?, Neural Computation 27 (2015) 1857–1871. doi:10/f7n8qq.

[54] P. A. Valdes-Sosa, Spatio-temporal autoregressive models defined over brain manifolds, Neuroinformatics 2 (2004) 239–250. doi:10/fs5xbw.

[55] J. Casorso, X. Kong, W. Chi, D. Van De Ville, B. T. Yeo, R. Liégeois, Dynamic mode decomposition of resting-state and task fMRI, NeuroImage 194 (2019) 42–54. doi:10/gfx53r.

[56] D. Bates, M. Mächler, B. Bolker, S. Walker, Fitting linear mixed-effects models using lme4, Journal of Statistical Software 67 (2015). doi:10/gcrnkw.

[57] E. Jolly, Pymer4: Connecting R and Python for linear mixed modeling, Journal of Open Source Software 3 (2018) 862. doi:10/gnzggv.

[58] L. Li, L. Zeng, Z.-J. Lin, M. Cazzell, H. Liu, Tutorial on use of intraclass correlation coefficients for assessing intertest reliability and its application in functional near-infrared spectroscopy–based brain imaging, Journal of Biomedical Optics 20 (2015) 050801. doi:10/gj7s8x.

[59] M. Rubinov, O. Sporns, Complex network measures of brain connectivity: Uses and interpretations, NeuroImage 52 (2010) 1059–1069. doi:10.1016/j.neuroimage.2009.10.003.

[60] M. E. J. Newman, Networks, second edition ed., Oxford University Press, Oxford, United Kingdom ; New York, NY, United States of America, 2018.

[61] J. D. Power, B. L. Schlaggar, C. N. Lessov-Schlaggar, S. E. Petersen, Evidence for hubs in human functional brain networks, Neuron 79 (2013) 798–813. doi:10.1016/j.neuron.2013.07.035.

[62] M. Pedersen, A. Omidvarnia, J. M. Shine, G. D. Jackson, A. Zalesky, Reducing the influence of intramodular connectivity in participation coefficient, Network Neuroscience 4 (2020) 416–431. doi:10.1162/netn_a_00127.

[63] R. Guimerà, L. A. Nunes Amaral, Functional cartography of complex metabolic networks, Nature 433 (2005) 895–900. doi:10.1038/nature03288.

[64] F. Klimm, J. Borge-Holthoefer, N. Wessel, J. Kurths, Gorka Zamora-López, Individual node’s contribution to the mesoscale of complex networks, New Journal of Physics 16 (2014) 125006. doi:10.1088/1367-2630/16/12/125006.

[65] S. Maslov, K. Sneppen, Specificity and stability in topology of protein networks, Science 296 (2002) 910–913. doi:10.1126/science.1065103.

[66] J. L. Ji, M. Spronk, K. Kulkarni, G. Repovš, A. Anticevic, M. W. Cole, Mapping the human brain’s cortical- subcortical functional network organization, NeuroImage 185 (2019) 35–57. doi:10.1016/j.neuroimage.2018.10.006.

[67] J. Li, R. Kong, R. Liégeois, C. Orban, Y. Tan, N. Sun, A. J. Holmes, M. R. Sabuncu, T. Ge, B. T. Yeo, Global signal regression strengthens association between resting-state functional connectivity and behavior, NeuroImage 196 (2019) 126–141. doi:10/gj8p69.

[68] R. Kashyap, R. Kong, S. Bhattacharjee, J. Li, J. Zhou, B. Thomas Yeo, Individual-specific fMRI-Subspaces improve functional connectivity prediction of behavior, NeuroImage 189 (2019) 804–812. doi:10/gft3tt.

[69] T. Ge, M. Reuter, A. M. Winkler, A. J. Holmes, P. H. Lee, L. S. Tirrell, J. L. Roffman, R. L. Buckner, J. W. Smoller, M. R. Sabuncu, Multidimensional heritability analysis of neuroanatomical shape, Nature Communications 7 (2016) 13291. doi:10/f9b8cv.

[70] D. J. Stekhoven, P. Buhlmann, MissForest–non-parametric missing value imputation for mixed-type data, Bioinformatics 28 (2012) 112–118. doi:10/dhxth8.

[71] T. Ge, T. E. Nichols, P. H. Lee, A. J. Holmes, J. L. Roffman, R. L. Buckner, M. R. Sabuncu, J. W. Smoller, Massively expedited genome-wide heritability analysis (MEGHA), Proceedings of the National Academy of Sciences 112 (2015) 2479–2484. doi:10/f63g67.

[72] A. C. Rencher, Methods of Multivariate Analysis, Wiley Series in Probability and Mathematical Statistics, 2nd ed ed., J. Wiley, New York, 2002.

[73] M. Helmer, S. Warrington, A.-R. Mohammadi-Nejad, J. L. Ji, A. Howell, B. Rosand, A. Anticevic, S. N. Sotiropoulos, J. D. Murray, On stability of Canonical Correlation Analysis and Partial Least Squares with application to brain-behavior associations, 2020. doi:10.1101/2020.08.25.265546.

[74] A. M. Winkler, O. Renaud, S. M. Smith, T. E. Nichols, Permutation inference for canonical correlation analysis, NeuroImage 220 (2020) 117065. doi:10.1016/j.neuroimage.2020.117065.

[75] H.-T. Wang, J. Smallwood, J. Mourao-Miranda, C. H. Xia, T. D. Satterthwaite, D. S. Bassett, D. Bzdok, Finding the needle in a high-dimensional haystack: Canonical correlation analysis for neuroscientists, NeuroImage 216 (2020) 116745. doi:10.1016/j.neuroimage.2020.116745.

[76] X. Zhuang, Z. Yang, D. Cordes, A technical review of canonical correlation analysis for neuroscience applications, Human Brain Mapping 41 (2020) 3807–3833. doi:10.1002/hbm.25090.

[77] S. M. Smith, T. E. Nichols, D. Vidaurre, A. M. Winkler, T. E. J. Behrens, M. F. Glasser, K. Ugurbil, D. M. Barch, D. C. Van Essen, K. L. Miller, A positive-negative mode of population covariation links brain connectivity, demographics and behavior, Nature Neuroscience 18 (2015) 1565–1567. doi:10.1038/nn.4125.

[78] A. Mihalik, J. Chapman, R. A. Adams, N. R. Winter, F. S. Ferreira, J. Shawe-Taylor, J. Mouraõ-Miranda, Canonical correlation analysis and partial least squares for identifying brain–behavior associations: A tutorial and a comparative study, Biological Psychiatry: Cognitive Neuroscience and Neuroimaging 7 (2022) 1055–1067. doi:10.1016/j.bpsc.2022.07.012.

[79] A. Schaefer, R. Kong, E. M. Gordon, T. O. Laumann, X.-N. Zuo, A. J. Holmes, S. B. Eickhoff, B. T. T. Yeo, Localglobal parcellation of the human cerebral cortex from intrinsic functional connectivity MRI, Cerebral Cortex 28 (2018) 3095–3114. doi:10/gd738m.

[80] E. B. Erhardt, E. A. Allen, Y. Wei, T. Eichele, V. D. Calhoun, SimTB, a simulation toolbox for fMRI data under a model of spatiotemporal separability, NeuroImage 59 (2012) 4160–4167. doi:10/cr4g9g.

[81] X.-N. Zuo, R. Ehmke, M. Mennes, D. Imperati, F. X. Castellanos, O. Sporns, M. P. Milham, Network centrality in the human functional connectome, Cerebral Cortex 22 (2012) 1862–1875. doi:10.1093/cercor/bhr269.

[82] M. Chong, C. Bhushan, A. Joshi, S. Choi, J. Haldar, D. Shattuck, R. Spreng, R. Leahy, Individual parcellation of resting fMRI with a group functional connectivity prior, NeuroImage 156 (2017) 87–100. doi:10.1016/j.neuroimage.2017.04.054.

[83] T. O. Laumann, A. Z. Snyder, Brain activity is not only for thinking, Current Opinion in Behavioral Sciences 40 (2021) 130–136. doi:10.1016/j.cobeha.2021.04.002.

[84] J. R. Cohen, The behavioral and cognitive relevance of time-varying, dynamic changes in functional connectivity, NeuroImage 180 (2018) 515–525. doi:10.1016/j.neuroimage.2017.09.036.

[85] E. A. Allen, E. Damaraju, S. M. Plis, E. B. Erhardt, T. Eichele, V. D. Calhoun, Tracking whole-brain connectivity dynamics in the resting state, Cerebral Cortex 24 (2014) 663–676. doi:10.1093/cercor/bhs352.

[86] D. Vidaurre, A. Llera, S. Smith, M. Woolrich, Behavioural relevance of spontaneous, transient brain network interactions in fMRI, NeuroImage 229 (2021) 117713. doi:10.1016/j.neuroimage.2020.117713.

[87] A. Eichenbaum, I. Pappas, D. Lurie, J. R. Cohen, M. D’Esposito, Differential contributions of static and timevarying functional connectivity to human behavior, Network Neuroscience 5 (2021) 145–165. doi:10.1162/netn_a_00172.

[88] A. S. Choe, M. B. Nebel, A. D. Barber, J. R. Cohen, Y. Xu, J. J. Pekar, B. Caffo, M. A. Lindquist, Comparing testretest reliability of dynamic functional connectivity methods, NeuroImage 158 (2017) 155–175. doi:10.1016/j.neuroimage.2017.07.005.

[89] W. H. Thompson, P. Fransson, The mean–variance relationship reveals two possible strategies for dynamic brain connectivity analysis in fMRI, Frontiers in Human Neuroscience 9 (2015). doi:10.3389/fnhum.2015.00398.

[90] C. Zhang, S. A. Baum, V. R. Adduru, B. B. Biswal, A. M. Michael, Test-retest reliability of dynamic functional connectivity in resting state fmri, NeuroImage 183 (2018) 907–918. doi:10.1016/j.neuroimage.2018.08.021.

[91] H. Jia, X. Hu, G. Deshpande, Behavioral relevance of the dynamics of the functional brain connectome, Brain Connectivity 4 (2014) 741–759. doi:10.1089/brain.2014.0300.

[92] B. Rashid, E. Damaraju, G. D. Pearlson, V. D. Calhoun, Dynamic connectivity states estimated from resting fMRI Identify differences among Schizophrenia, bipolar disorder, and healthy control subjects, Frontiers in Human Neuroscience 8 (2014). doi:10.3389/fnhum.2014.00897.

[93] S. Marek, B. Tervo-Clemmens, F. J. Calabro, D. F. Montez, B. P. Kay, A. S. Hatoum, M. R. Donohue, W. Foran, R. L. Miller, T. J. Hendrickson, S. M. Malone, S. Kandala, E. Feczko, O. Miranda-Dominguez, A. M. Graham, E. A. Earl, A. J. Perrone, M. Cordova, O. Doyle, L. A. Moore, G. M. Conan, J. Uriarte, K. Snider, B. J. Lynch, J. C. Wilgenbusch, T. Pengo, A. Tam, J. Chen, D. J. Newbold, A. Zheng, N. A. Seider, A. N. Van, A. Metoki, R. J. Chauvin, T. O. Laumann, D. J. Greene, S. E. Petersen, H. Garavan, W. K. Thompson, T. E. Nichols, B. T. T. Yeo, D. M. Barch, B. Luna, D. A. Fair, N. U. F. Dosenbach, Reproducible brain-wide association studies require thousands of individuals, Nature 603 (2022) 654–660. doi:10.1038/s41586-022-04492-9.

[94] C. T. Ellis, C. Baldassano, A. C. Schapiro, M. B. Cai, J. D. Cohen, Facilitating open-science with realistic fMRI simulation: Validation and application, PeerJ 8 (2020) e8564. doi:10/ght935.

[95] J. E. Chen, J. R. Polimeni, S. Bollmann, G. H. Glover, On the analysis of rapidly sampled fMRI data, NeuroImage 188 (2019) 807–820. doi:10/gfvhhv.

[96] S. Bollmann, A. M. Puckett, R. Cunnington, M. Barth, Serial correlations in single-subject fMRI with sub-second TR, NeuroImage 166 (2018) 152–166. doi:10/gcr9cx.

[97] D. Rangaprakash, G.-R. Wu, D. Marinazzo, X. Hu, G. Deshpande, Hemodynamic response function (HRF) variability confounds resting-state fMRI functional connectivity, Magnetic Resonance in Medicine 80 (2018) 1697– 1713. doi:10/gkzm4c.

[98] E. Tagliazucchi, H. Laufs, Multimodal imaging of dynamic functional connectivity, Frontiers in Neurology 6 (2015). doi:10.3389/fneur.2015.00010.

[99] M. X. Cohen, Analyzing Neural Time Series Data: Theory and Practice, Issues in Clinical and Cognitive Neuropsychology, The MIT Press, Cambridge, Massachusetts, 2014.

